# Bayesian copy number detection and association in large-scale studies

**DOI:** 10.1101/2020.01.24.918672

**Authors:** Stephen Cristiano, David McKean, Jacob Carey, Paige Bracci, Paul Brennan, Michael Chou, Mengmeng Du, Steven Gallinger, Michael G. Goggins, Manal Hassan, Rayjean Hung, Robert Kurtz, Donghui Li, Lingeng Lu, Rachel Neale, Sara Olson, Gloria Petersen, Kari Rabe, Jack Fu, Harvey Risch, Gary Rosner, Ingo Ruczinski, Alison P. Klein, Robert B. Scharpf

## Abstract

Germline copy number variants (CNVs) increase risk for many diseases, yet detection of CNVs and quantifying their contribution to disease risk in large-scale studies is challenging. We developed an approach called CNPBayes to identify latent batch effects, to provide probabilistic estimates of integer copy number across the estimated batches, and to fully integrate the copy number uncertainty in the association model for disease. We demonstrate this approach in a Pancreatic Cancer Case Control study of 7,598 participants where the major sources of technical variation were not captured by study site and varied across the genome. Candidate associations aided by this approach include deletions of 8q24 near regulatory elements of the tumor oncogene *MYC* and of Tumor Supressor Candidate 3 (*TUSC3*). This study provides a robust Bayesian inferential framework for estimating copy number and evaluating the role of copy number in heritable diseases.

## Introduction

Germline copy number variants (CNVs) can be identified from hybridization-based arrays and capture-based sequencing with measures of abundance derived from intensities and normalized read depth, respectively. Biological and technical sources of heterogeneity of these measurements are intricately related. For example, the GC composition of genomic DNA effects PCR efficiency and leads to auto-correlated measures of DNA abundance across the genome [1–4]. These effects have been shown to be heterogeneous across the genome and to differ in both magnitude and direction between samples [1, 5, 6]. Hidden Markov models and nonparametric segmentation algorithms for CNV detection over-segment low-quality data where these effects are the most pronounced, contributing to false positive deletion and duplication calls.

For studies with hundreds to thousands of samples, estimation of copy number at regions known to harbor CNVs has the potential to improve sensitivity and specificity as technical sources of variation across the genome are largely controlled when limited to a focal genomic region (less than 1 MB) and variation between samples can be explicitly modeled [7–12]. Such CNV regions are of particular interest for a comprehensive assessment of common genetic variants and their relationship to disease. However, scaling marginal models to CNV regions across the genome and to thousands of samples has proved challenging. The sources of technical variation giving rise to batch effects are typically unknown. Standard approaches for estimating latent batch effects in high-throughput experiments such as surrogate variable analysis are not appropriate when the biological variation of interest (copy number) is also unknown [13]. In addition, the statistical framework for copy number estimation must flexibly accommodate deletions and duplications of varying size and allele frequencies. Symptomatic of the challenges in copy number analyses and the limitations of current methods, genome-wide association studies rarely incorporate copy number in the initial publication despite their well established role in neurodevelopmental disorders [14–16] and cancer [17]. Previous genome-wide studies of pancreatic cancer and copy number have been limited in size with fewer than 250 pancreatic cancer patients [18, 19].

Here, we performed genome-wide copy number analysis for 3,974 cases and 3,624 controls in PanC4 using Illumina’s OmniExpress Exome array. We developed methods for identifying latent batch effects at CNV regions from commonly available experimental data on the samples. The effects of copy number and batch on measures of DNA abundance were modeled hierarchically through implementation of Bayesian mixture models. For the association model, we used Markov Chain Monte Carlo (MCMC) to incorporate the uncertainty of the integer copy numbers in a logistic regression model of pancreatic cancer risk.

## Results

### Overview of study

DNA specimens from 7,598 European ancestry participants in this consortium were collected at 9 study sites using varying methods of DNA extraction [20]. Randomization of samples to chemistry plates, DNA amplification by PCR, and SNP genotyping using Illumina’s OmniExpress Exome-8 array were performed centrally at the Center for Inherited Disease Research (CIDR) (Figure 1). CNV regions were extracted from the 1000 Genomes project [21] or identified from analysis of the PanC4 samples. Low-level copy number summaries were obtained for each participant by computing the median log_2_ *R* ratios across available markers from the Illumina array spanned by the candidate CNV region. Independently for each region, we identified latent batch effects in the low-level summaries and fit a Bayesian hierarchical mixture model across the estimated batches using CNPBayes. To model the relationship between copy number and pancreatic cancer risk, we fit a Bayesian logistic regression model that included integer copy number as a covariate measured with error. The copy number measurement error for each participant was obtained from the posterior probabilities in the CNPBayes hierarchical model.

**Figure 1:**
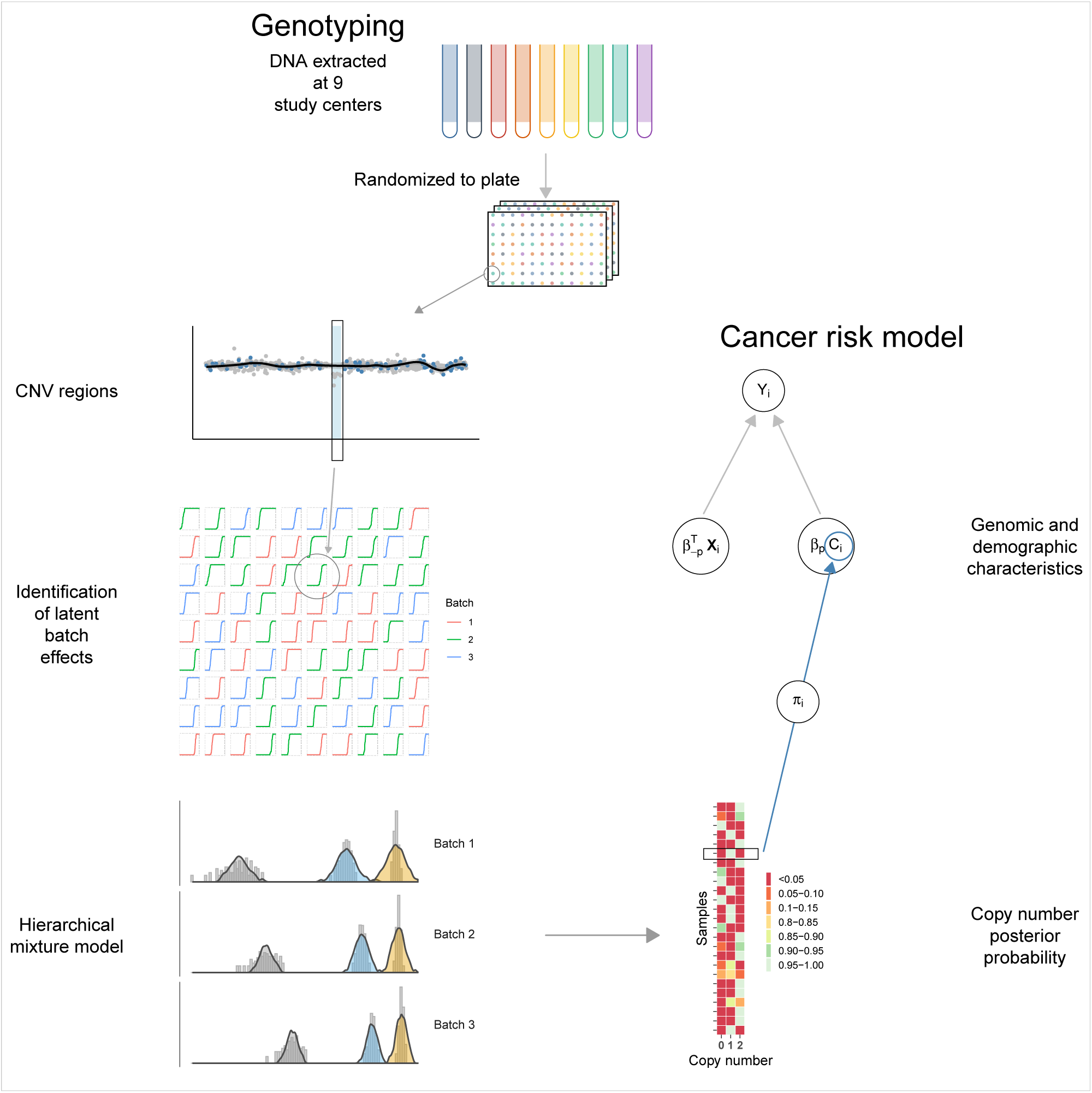
Overview of sample processing, estimation of batch effects and copy number, and risk model for pancreatic cancer. DNA samples for pancreatic cancer cases and healthy controls were obtained from 9 different study centers and processed centrally where samples were randomized to chemistry plates. Initial preprocessing of these samples identified candidate CNV regions. As the principal sources of batch effects were unknown, we developed an approach to identify latent batch effects and to genotype these samples via a Bayesian hierarchical mixture model. Uncertainty of the copy number genotypes was propagated from the genomic analyses to the Bayesian logistic regression model for pancreatic cancer risk.

### Copy number analyses

Log_2_*R* ratios for each participant were GC corrected using loess. Measures of data quality following GC-correction include the median absolute deviation and lag-10 autocorrelation of autosomal log_2_ *R* ratios ordered by genomic position. Data quality was high for the majority of PanC4 participants (Figure S1A), though approximately 11% of participants had autocorrelations greater than 0.1. To reduce the spatial autocorrelation, we developed a scatterplot smoother for the log_2_ *R* ratios that was locally weighted by genomic position (Methods). Following the spatial correction, nearly all samples (≈ 98%) had low autocorrelation (Figure S1B). Rare and common CNVs identified by a 5-state hidden Markov model before and after spatial correction revealed near perfect concordance for samples in the first nine deciles of autocorrelation (high quality samples) with sharply lower concordance among samples in the highest decile irrespective of CNV size (Figure S2).

To evaluate whether copy number inference could be improved by multi-sample methods that directly incorporate batch and other technical sources of variation between samples, we focused our analysis on 217 regions from the 1000 Genomes Project where CNVs were reported in at least 0.1% of European ancestry participants and that encompassed at least four probes on the Illumina OmniExome array (Table S1). Additionally, we identified 46 regions for which deletions or duplications were identified in at least 2% of the PanC4 participants by the hidden Markov model applied to the spatially corrected log_2_ *R* ratios. Collectively, the 263 regions comprised 11.5 Mb of the coding genome and 6.4 Mb of the non-coding genome.

Available multi-sample methods for modeling copy number assume the major sources of batch effects are known (e.g., laboratory or study site). Here, DNA samples were collected from multiple study sites and processed on 94 chemistry plates at a central lab. To identify batch surrogates for the central lab, we developed an approach for grouping chemistry plates with a similar median log_2_ *R* ratio empirical cumulative distribution function (eCDF) (Figures 2A and 2B). As an example of these sources of heterogeneity at a single CNV region on chromosome 4, we summarized the log_2_ *R* ratios for 6,026 high quality samples by the first principal component (PC1) and stratified the PC1 summaries by study site (Figure 3A) or PCR batch surrogates (Figure 3B). While the density of PC1 is bimodal when stratified by study site and consistent with a copy number polymorphism, stratification by the eCDF-derived batch surrogates revealed obvious batch effects (e.g., plate group C with 567 samples and plate group E with 786 samples; Figure 3B). As PCR efficiency is known to be affected by GC content and can vary along the genome, we identified batch surrogates for each CNV region. The median number of batches across the 263 CNV regions was 4 with multiple batches identified for the majority of regions (Figure S3).

**Figure 2:**
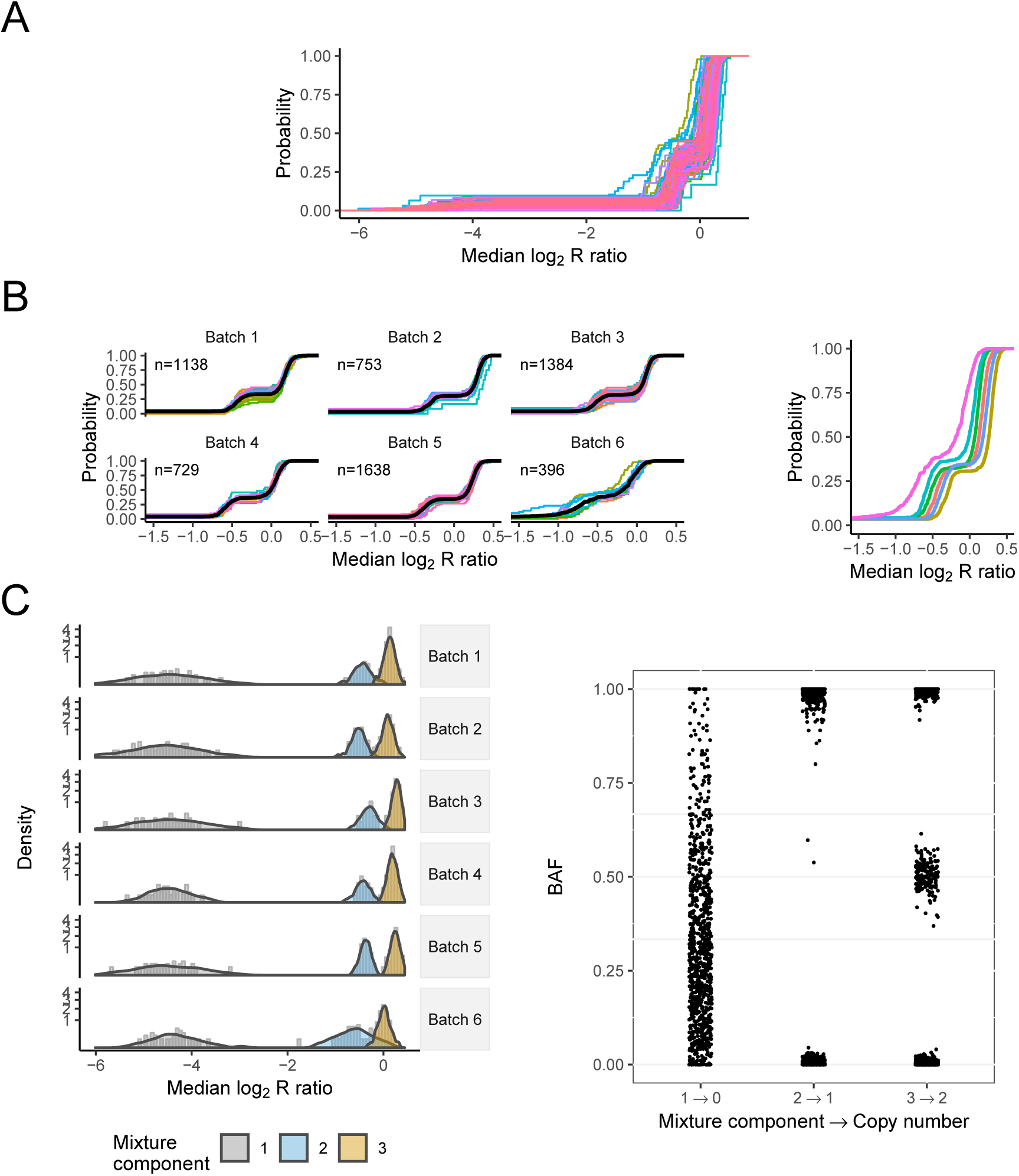
Identification of batch surrogates. **(A)** Plate-specific empirical cumulative distribution functions (eCDFs) of the average log_2_ *R* ratio for a region on chr5 (155,475,886-155,488,649bp). **(B)** The plate-specific eCDFs were grouped by Kolmogorov-Smirnov test statistics, forming batches. The batch-specific eCDFs after grouping plates (right). The eCDFs between batches typically differed by a location shift, though here Batch 6 also captured samples with higher variance. **(C)** Single- and multi-batch mixture models were evaluated at each CNP and ranked by the marginal likelihood. Densities from the posterior predictive distributions overlay the histograms of the 3-component multi-batch model (left). Adjusted for batch, only three components were needed to fit the apparent deletion polymorphism. B allele frequencies were used to genotype the mixture components (right).

**Figure 3:**
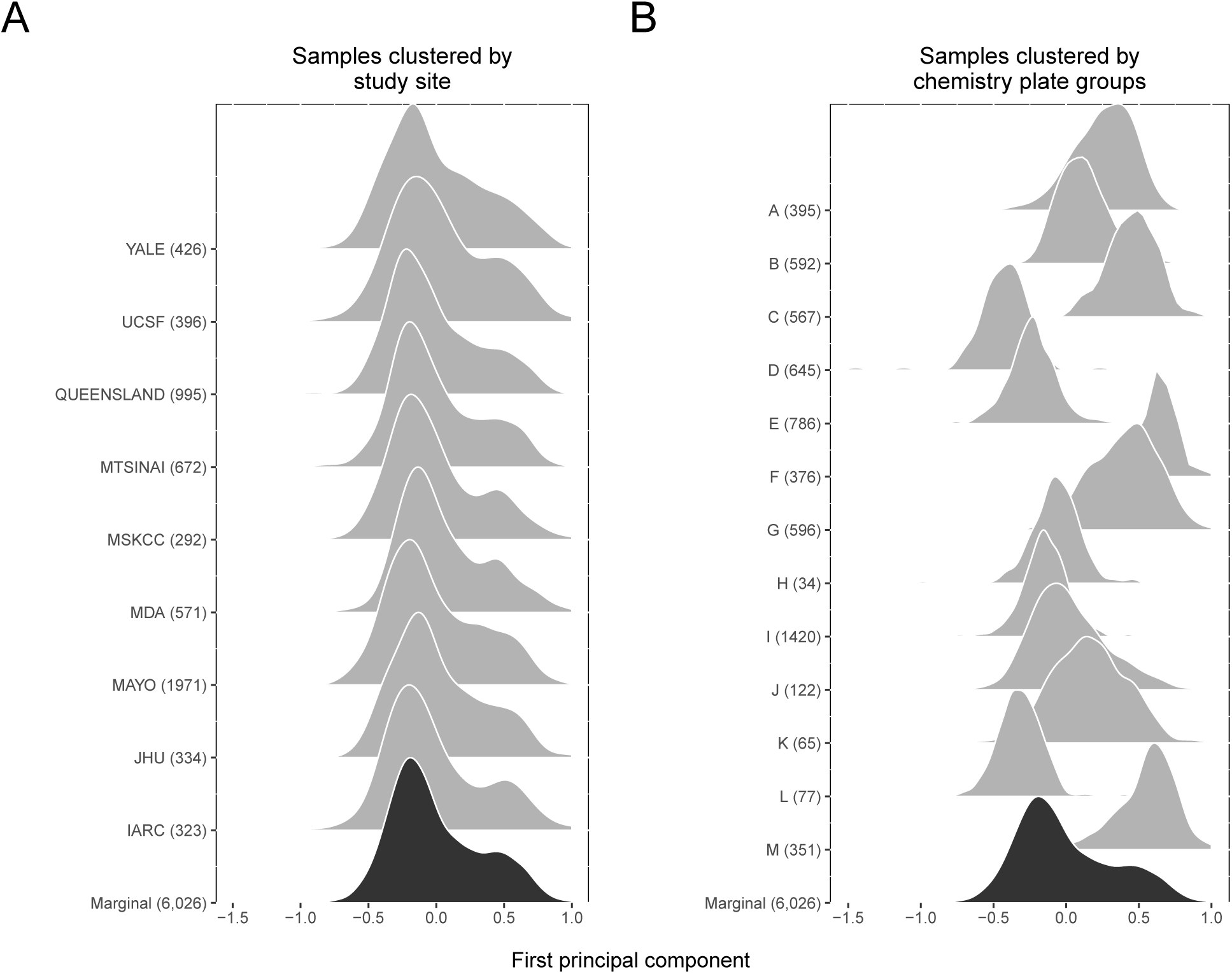
Study site does not capture the major sources of technical variation. Hybridization intensities were available for four probes in a CNP region on chr4 spanning 9,370,866 bp - 9,410,140 bp (CNP_051). Restricting our analysis to high quality samples, we used the first principal component (PC1) as a one-dimensional summary of the 4 × 6,026 matrix of log_2_ *R* ratios. The density of the PC1 summaries marginally (black) and stratified by study site (gray) are bimodal, suggesting a copy number polymorphism **(A)**. However, stratification of the PC1 summaries by grouping chemistry plates with similar eCDFs reveals an obvious batch effect **(B)**. For example, chemistry plates in group E comprised of 786 samples originating from all nine study sites has a markedly different distribution than the 567 samples processed on group C chemistry plates.

Our sampling model for the median log_2_ *R* ratio is a mixture of *t* distributions with component-specific means and variances modeled hierarchically across batches (Figure 2C). To overcome challenges inherent to modeling rare homozygous deletions when these were apparent in a small number of samples, we augmented the observed data with simulated deletions. For selecting the number of mixture components, CNPBayes provides an option to estimate the marginal likelihood through additional MCMC simulations [22] and the posterior predictive distribution to aid assessments of goodness of fit. Following the probabilistic assignment of samples to mixture components, we genotyped the mixture components using the available B allele frequencies (BAFs) at SNPs (Figure 2C). From the 263 CNV regions, 25 regions contained samples with duplications, 132 regions contained samples with deletions, and 24 regions contained samples with deletions as well as samples with duplications. Allele frequencies from the genotyped duplications and deletions in the PanC4 controls were consistent with percentages reported in the 1000 Genomes Project. We identified a median of 17 additional CNVs per sample by the mixture model that were not identified by the hidden Markov model (Figures S4 and S5). On average, CNVs spanned 6 SNPs (IQR: 5-8) and were 12.6kb in size (IQR: 10.9kb - 17.6kb).

### Simulation

To benchmark the sensitivity and specificity of this approach when the true genotypes were known, we extracted high quality data from a subset of HapMap phase III samples (n = 990) processed on 16 chemistry plates and hybridized to Affymetrix 6.0 chips. A 109 kb region on chr 4 containing 1 SNP and 53 nonpolymorphic markers spans a deletion polymorphism with an allele frequency near 22%. We increased the level of difficulty for genotyping these samples by increasing the variance and/or shifting the location of the probe-level data in a subset of the chemistry plates. For each simulated dataset, we fit both `CNVCALL`, a mixture model developed for the analysis of the Welcome Trust Case Control Consortium, and CNPBayes [9, 23]. While we did not provide the true batch labels to either method, CNPBayes estimated the batches from the plate surrogates. With no simulated batch effects, CNPBayes and `CNVCALL` had nearly identical performance with near perfect sensitivity and specificity (area under the receiver operator characteristic curve (AUC) > 0.99). However, for simulated datasets with batch effects in the mean or variance, accuracy of `CNVCALL` decreased by an average of 25% while performance characteristics of CNPBayes remained qualitatively similar (Figure S6).

### Risk model for pancreatic cancer

To evaluate whether changes in germline copy number effect pancreatic cancer risk, we fit a Bayesian logistic regression model at each CNV region. Uncertainty of the copy number assignment for each participant was incorporated in the regression model by sampling the integer copy number from a multinomial distribution parametrized by posterior probabilities from CNPBayes at each scan of the MCMC. As case-control status was unevenly distributed between the high and low data quality sample collections 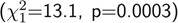, the regression model included an interaction between copy number and data quality (Methods) as well as a single binary parameter *z*_*c*_ multiplying both of these terms that allows the slopes to be exactly zero. The posterior mean of *z*_*c*_ provides an estimate of the probability of an association with copy number (Figure 4 and Table S2). Additional covariates included age, gender, and the first three principal components previously of the SNP genotypes [20].

**Figure 4:**
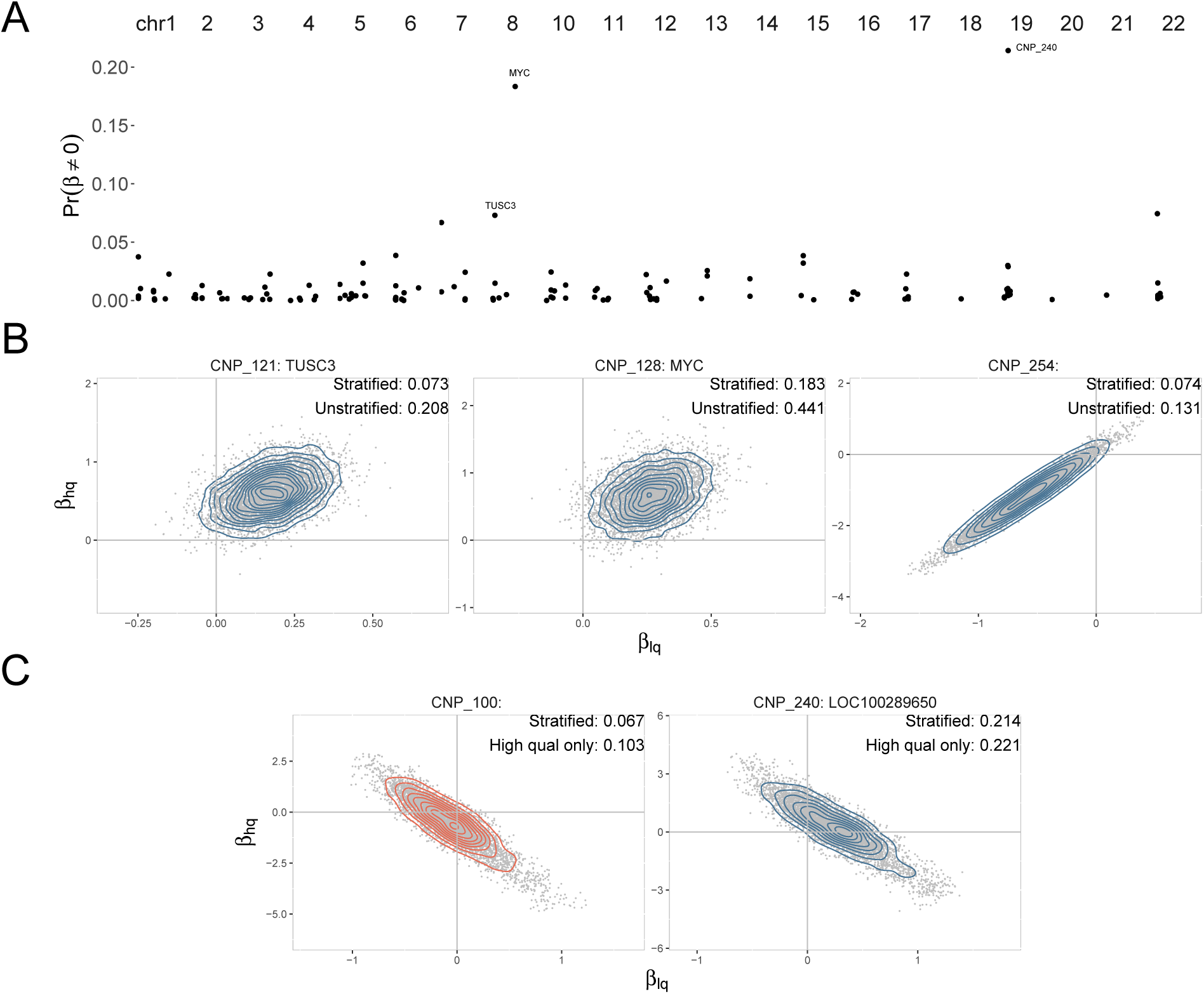
Bayesian regression models for pancreatic cancer risk. To incorporate uncertainty of the copy number assignment from the low-level data, the integer copy number was sampled from the subject-specific posterior probabilities provided by CNPBayes at each iteration of the MCMC. While batch effects on CNV inference were already accounted for in the low and high quality sample collections, an imbalance of the pancreatic cancer cases between these collections warranted a stratified model with an interaction between copy number and data quality and an indicator, *z*_*c*_, multiplying these coefficients that allowed the slopes to be exactly zero. For CNV regions where copy number inference was unaffected by data quality and associated with pancreatic cancer risk, regression coefficients for the low and high quality collections were positively correlated and the posterior mean of *z*_*c*_ (upper right corner) increased in the more powerful unstratified analysis using all 7,598 samples **(B)**. By contrast, negatively correlated coefficients indicated an effect of data quality on CNV inference confirmed by visual inspection and the appropriate follow-up analysis and estimated probability of association was limited to the high quality sample collection **(C)**.

Genome-wide posterior probabilities of association between copy number and pancreatic cancer risk were near zero for most CNV regions (Figure 4A). Using the posterior probability of association to select CNV regions that may modulate pancreatic cancer risk, we characterized the joint distribution of the regression coefficients for the high and low quality samples under simulations for which *z*_*c*_ was one (Figures 4B and 4C)). Participants with two copies of the Tumor Supressor Candidate 3, *TUSC3*, had a 20% increased odds of pancreatic cancer compared to individuals with germline hemizygous deletions in this gene (90% credible interval (CI) for odds ratio: 1.01 - 1.39). While the direction of this effect is inconsistent with its putative role as a tumor suppressor, up-regulation of *TUSC3* and possible oncogenic roles for this gene have been reported in cancers including non-small cell lung cancer, colorectal, thyroid, and head and neck cancers [24–27]. Among non-coding regions, we found that deletions for a CNV region in 8q24 were associated with a reduced risk of pancreatic cancer (90% CI: 1.09-1.59). Chromosome 8q24 has been implicated in many cancers and is known to contain regulatory elements for the tumor oncogene *MYC* located at 128,748,315-128,753,680 bp [28]. We have previously demonstrated the association of SNPs in this region with an increased risk of pancreatic cancer [29, 30]. As copy number regression coefficients at CNV regions spanning *TUSC3* and near *MYC* were positive and highly correlated for both the low and high quality sample collections, an unstratified analysis using all 7,598 participants for these genes is potentially more powerful and doubled the posterior probability of association for these genes (Figure 4B). Overall, our approach provides conservative measures of the association between copy number and pancreatic cancer risk across the genome, accounting for latent batch effects and copy number uncertainty separately for samples where data quality was more compromised.

## Discussion

We performed a genome-wide analysis of germline copy number variants in the largest study to date of pancreatic cancer, implementing approaches to correct for latent batch effects and risk models that incorporate uncertainty of the copy number estimates. As the batch effects we identified were likely related to differences in PCR efficiencies that can vary across the genome and between groups of samples processed on different chemistry plates within a single laboratory (not between study sites), we identified and adjusted for batch effects in a region-dependent manner in contrast to alternative methods. Using this approach, nearly 70% of CNV regions analyzed had multiple batches that were related to chemistry plates and not the individual laboratories that contributed samples. These batch effects are not limited to hybridization-based arrays and are present in any technology relying on PCR amplification including most capture-based sequencing protocols. As studies become increasingly large-scale with inevitable batch effects and heterogeneity in sample quality, the scalable approach provided by CNPBayes will be helpful for modeling unwanted technical variation and avoiding the potential confounding between batch effects and copy number when evaluating disease risk.

Using the methods outlined in this study and our a priori probability that the regression coefficients for many copy number alterations will not be associated with cancer risk, only germline deletions of *TUSC3* and near *MYC* had an appreciable posterior probability of an association with pancreatic cancer risk. An increased prevalance of germline deletions at *TUSC3* and *MYC* among healthy participants has not been previouly implicated in pancreatic cancer, though upregulation of expression of these genes has been implicated in some cancers in an apparent tissue-dependent manner.

While we evaluated copy number at both known and HMM-discoverable CNV regions for pancreatic cancer risk, more sensitive technologies for identifying smaller CNV regions with potentially rare germline CNVs among cancer patients are needed. Whether moscaic copy number alterations in hematopoietic cells could further modulate risk has not been evaluated [31–34]. Finally, we assumed an additive model for integer copy number and the log odds of cancer risk. Dominant and recessive mechanisms of genotype-phenotype associations are possible and the evidence for these models using Bayes factors could be averaged with weights reflecting our a priori beliefs.

## Methods

### The Pancreatic Cancer Case and Control Consortium

Clinical and demographic characteristics of the cases and controls in PanC4 and recruitment methods have been previously described [20]. All samples were processed using GenomeStudio (version 2011.1, Genotyping Module 1.9.4). For GC-correction, we sampled a random subset of 30,000 Illumina probes, fit LOESS with span 1/3 to the scatterplot of log_2_ *R* ratios and probe GC content, and predicted the log_2_ *R* ratios for the full probeset from the LOESS model. For spatial correction, we applied LOESS to the GC-corrected log_2_ *R* ratios at SNPs with balanced allele frequencies (0.4 < *BAF* < 0.6) ordered by genomic position within each chromosome arm and predicted the GC-corrected log_2_ *R* ratios for the full probeset, including SNPs with imbalanced allele frequencies. The residuals from the spatial LOESS were used in all downstream analyses with CNPBayes.

### CNV regions

CNV regions identified for further analysis by CNPBayes were obtained from the collection of CNVs identified from a hidden Markov model as well as known CNV regions from the 1000 Genomes Project. For the former, we fit a 5-state hidden Markov model implemented in the R package VanillaICE (version 1.40.0) using default parameter settings [35]. To obtain a high confidence call set, we removed CNVs with fewer than 10 probes, CNVs with posterior probability less than 0.9, and restricted inference to autosomal chromosomes. To assess the effect of spatial adjustment on copy number inference, we stratified the samples into deciles of median absolute deviation and autocorrelation and compared the results of the 5-state hidden Markov model fit after GC-correction to the CNVs identified after spatial correction. Concordance of CNVs identified by the hidden Markov models was defined by ≥ 50% reciprocal overlap [36].

CNV regions were defined by the set of non-overlapping disjoint intervals across the pooled set of all CNVs from cases and controls. We computed the number of subjects with a CNV overlapping each disjoint interval, retaining intervals where CNVs were identified in at least 150 participants. Among the disjoint intervals, we defined the CNV region as the minimum start and maximum end for which at least 50 percent of the copy number altered samples had a CNV. For CNV regions obtained from the 1000 Genomes Project, we excluded regions that did not span at least 4 markers on the OmniExpress array.

### Batch effects

We evaluated both chemistry plate and DNA extraction method as surrogates for batch effects. With provisionally defined batches, we compared the eCDF of the mean *r* between two batches (excluding samples with log_2_ *R* ratio < −1) by the Kolmogorov-Smirnov (K-S) test statistic. For two batches without a statistically significant difference in the K-S statistic at a type 1 error of 0.01, the batches were combined into a single new batch. This procedure was applied recursively at each CNV region until no further grouping of batch surrogates could be obtained.

### Hierarchical Bayesian mixture model

Hierarchical Bayesian mixtures of *t*-distributions were used to cluster median log_2_ *R* ratios within a CNV region. Let *r*_*ib*_ and *z*_*ib*_ denote the observed one-dimensional summary of log_2_ ratios measured from the array and the true (but latent) mixture component, respectively, for the *i*th individual in batch *b*. Given *z*_*ib*_ is some integer *h* (*h* ∈ {1, …, *K*} for a *K*-component model), our sampling model for the observed data is a shifted and scaled *t*-distribution with *d* degrees of freedom, mean *θ*_*hb*_, and standard deviation *σ*_*hb*_ that depends on batch:

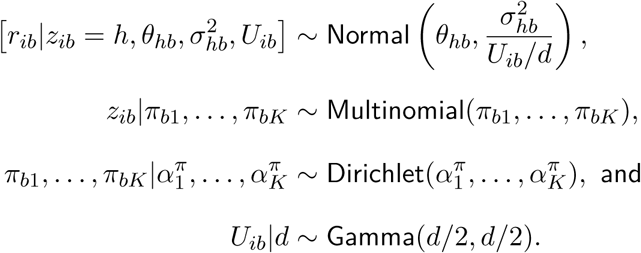

The degrees of freedom *d* controls robustness to outliers with larger values approximating a mixture of normal distributions [37]. To stabilize the mean and precision of batches having few samples, we model these parameters hierarchically with computationally convenient conjugate priors. Our sampling model for the batch means is normal and the precisions are Gamma,

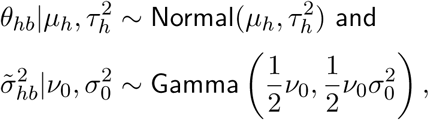

with *µ*_*h*_ representing the overall mean for component *h, τ*_*h*_ capturing the heterogeneity of the batch-specific means, and 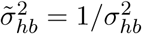. Conjugate priors on 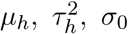, and *v*_0_ are given by

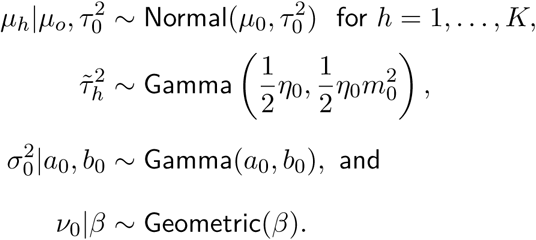

Label switching is well known in Bayesian mixture models. In addition to visual inspection of the chains, we compared the ordering of parameter means for subsequences of the chains. Label switching occurred most often when the number of components specified was too large and these models were discarded. In addition, we use an informative prior on 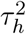 that governs the heterogeneity of the mean for mixture component *h* across the batches (Table S3). This prior discourages label switching at bona fide copy number polymorphisms since this would typically result in a large variance in component means.

As all priors were conjugate, we used Gibbs sampling to approximate the joint posterior distribution of 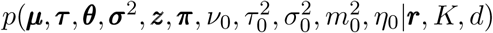. We refer to the above implementation of the Gibbs sampler with batch-specific means and variances as the multi-batch (MB) model. CNPBayes provides several more parsimonious alternatives to the MB model, including a pooled variance model (MBP) with a single variance estimate per batch. In addition, we evaluated models with a single batch (SB) and a single batch model with pooled variances (SBP) that are special cases of the MB and MBP models, respectively. Hyper-parameters used in the MB, MBP, SB, and SBP models were the defaults in version 1.11.2 of CNPBayes (Table S3).

### Implementation

Heavy-tailed marginal distributions of the one-dimensional log_2_ *R* ratio summaries were often a consequence of batch effects. When the latent batch effects were identified as previously described, normal mixtures performed well and CNPBayes was fit using *t* distributions with 100 degrees of freedom. For studies of germline CNVs such as PanC4, extreme observations in the left-tail correspond to homozygous deletions. To ensure these observations were not fit as outliers, we performed a data augmentation step where additional values were simulated based on the observed mean and standard deviation of the apparent homozygous deletions. For rare deletions, the data augmentation ensured that the hierarchical model appropriately borrowed information between batches in the event that some batches did not contain a sample with the homozygously deleted locus.

As fitting hierarchical Bayesian mixture models is computationally intensive, we implemented ad hoc procedures to reduce computation (see also Scalability and Software). First, we considered only 3 and 4 component models when homozygous deletions were apparent (one- and two-component models were not evaluated). Secondly, MB and MBP models were only evaluated when more than 2% of the samples had a posterior probability < 0.99 in the SB and/or SBP models. Thirdly, for each model under consideration, we independently initialized 10 models with parameters randomly sampled from their priors and ran a short burnin of 200 iterations for each model. Only the model with the largest log likelihood was selected for an additional 500 burnin simulations and 1000 simulations post-burnin. Finally, to aid comparison between hierarchical SB, SBP, MB, and MBP models, the CNPBayes package implements Chib’s method to estimate the marginal likelihood [22], allowing estimates of the relative evidence between two models by Bayes factors. However, as estimation of the marginal likelihoods requires additional MCMC simulations, we only computed marginal likelihoods when the difference of simple post-hoc statistical summaries, such as the log likelihood evaluated at the last iteration, was small (e.g., < 10). CNPBayes automatically provides posterior predictive distributions of the CNV region summaries for goodness of fit assessments, allowing the user to verify that the selected model is not simply the best of many poor fitting models.

### Genotyping mixture components

For genotyping the mixture components at a CNP region, our goal was to identify the set of integer copy numbers that would most likely give rise to the observed B allele frequencies at SNPs in this genomic region. We excluded samples that were not assigned to a single mixture component with high posterior probability since these would be less informative. Denoting the mapping of mixture component indices *h* to integer copy numbers by *f*(*h*), the likelihood across SNPs indexed by *j* and samples indexed by *I* conditional on the mapping is

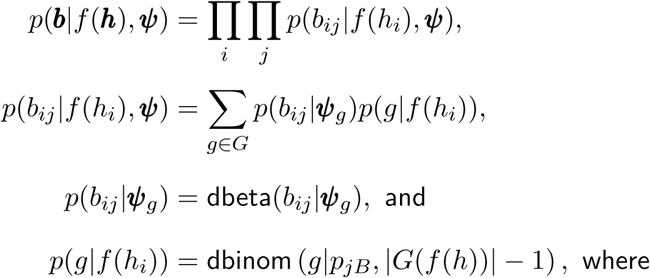

dbeta and dbinom are shorthand for the densities for the beta and binomial distributions. For the binomial density, |·| denotes the cardinality of the set and *p*_*jB*_ the frequency of the B allele at SNP *j* in the population of PanC4 participants. The above likelihood averages over the set *G* of possible allele specific copy numbers ordered by the number of B alleles and indexed by *g* (e.g., *G*(2) ∈ {*AA, AB, BB*}; Table S4). Shape and scale parameters (***ψ***_*g*_) for the Beta distribution conditional on the allelic copy numbers are provided in Table S5. Evaluating one-to-one (e.g., *f* ({1, 2, 3}) → {0, 1, 2} for a deletion polymorphism) and many-to-one mappings (e.g., *f* ({1, 2, 3}) → {2, 2, 2}), we selected the mapping that maximized the above likelihood on the log-scale.

### Simulation

Affymetrix 6.0 data for 990 phase 3 HapMap samples processed on 16 chemistry plates were obtained from Wellcome Sanger Institute (https://www.sanger.ac.uk/resources/downloads/human/hapmap3.html) [38]. A region on chr4 70,122,981-70,231,746 containing 53 nonpolymorphic markers and 1 SNP spans a common copy number polymorphism. To establish a baseline for which both CNPBayes and `CNVCALL` correctly identify the copy number for all samples, we subtracted 3 from the log_2_ *R* ratios for samples with apparent homozygous deletions. To simulate batch effects, we simulated a Bernoulli random variable with probability of success 0.5 for each of the 16 chemistry plates. For a plate *k* where the Bernoulli random deviate was 1, we rescaled the data by a factor *ξ* and shifted the means by a normal random deviate centered at *δ* such that the simulated log_2_ *R* ratio (*r*^*^) for marker *i* in sample *j* with integer copy number *c* becomes 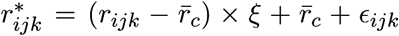, where *ϵ*_*ijk*_ ∼ *N* (*δ*, 0.02^2^) for values of *δ* ∈ {0, 0.3, 0.4, 0.5} and *ξ* ∈ {1, 1.25, 1.50, 1.75, 2}. Applying `CNVCALL` to this data, the matrix of ***r***^*^ was summarized by the first principal component and mixture models with 3-5 components were evaluated using default parameters. As `CNVCALL` merges mixture components based on the extent of overlap of the component-specific densities but does not genotype the merged mixture components, we subtracted one from the merged mixture component indices. For CNPBayes, we explored SB, SBP, MB, and MBP models of 3 - 4 components with chemistry plate as the surrogate variable, median 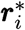 as one-dimensional summaries for each sample, and default values for hyperparameters. Mixture components were genotyped using the B allele frequencies from the SNP in this region as previously described.

#### Bayesian logistic regression model for pancreatic cancer

For each CNP region, we modeled the case-control status *y*_*i*_ for individual *i* as:

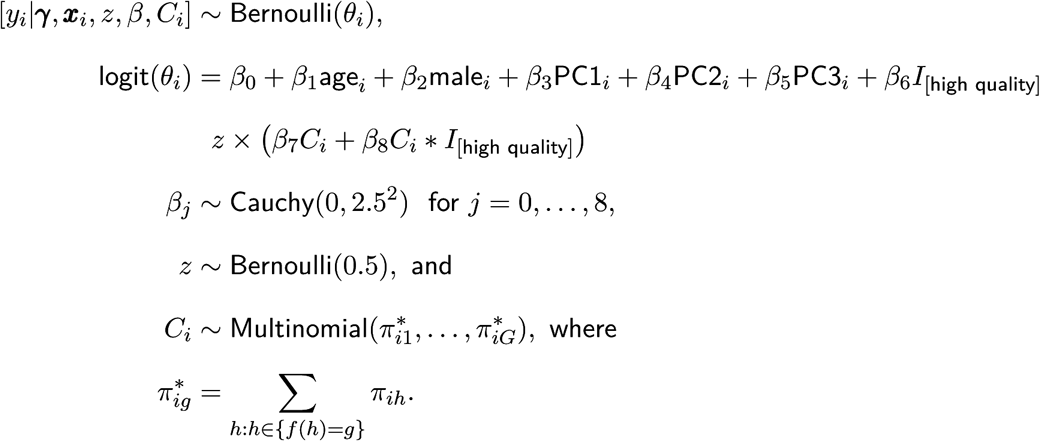

All continuous independent variables were mean centered, including PC1, PC2, and PC3 that denote the first three principal components of the SNP genotype matrix in PanC4 [20]. An indicator for the collection of high quality samples for CNV analyses, *I*_[high quality]_, was defined as 1 for samples in this set and 0 otherwise. As the integer copy number *C*_*i*_ was not observed, we treated *C*_*i*_ as a parameter measured with error given by the aggregated posterior probabilities of the mixture component indices after genotyping, 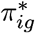. We used JAGS version 4.3.0 with a thin parameter of 25 and 5000 iterations to obtain posterior distributions of these parameters by MCMC [39].

### Scalability and software

Hierarchical mixture models were fit to a random sample of 1000 observations from the 7,598 available participants at each CNP region, and to all samples with apparent homozygous deletions. We parallelized our analysis so that all regions were evaluated simultaneously. Bayesian logistic regression models fit independently to each CNV region were also evaluated in parallel. CNPBayes is available from github (https://github.com/scristia/CNPBayes).

## Acknowledgements

We would like to thank Aravinda Chakravarti, Ann Oberg, Irene Orlow, and members of our laboratories for criticial review of this manuscript. This work was supported in part by the US National Institutes of Health grants 5R01CA154823, CA006973, and CA062924.

