## Supplemental Figures for "Bayesian copy number detection and association in large-scale studies"

### 1 Figures

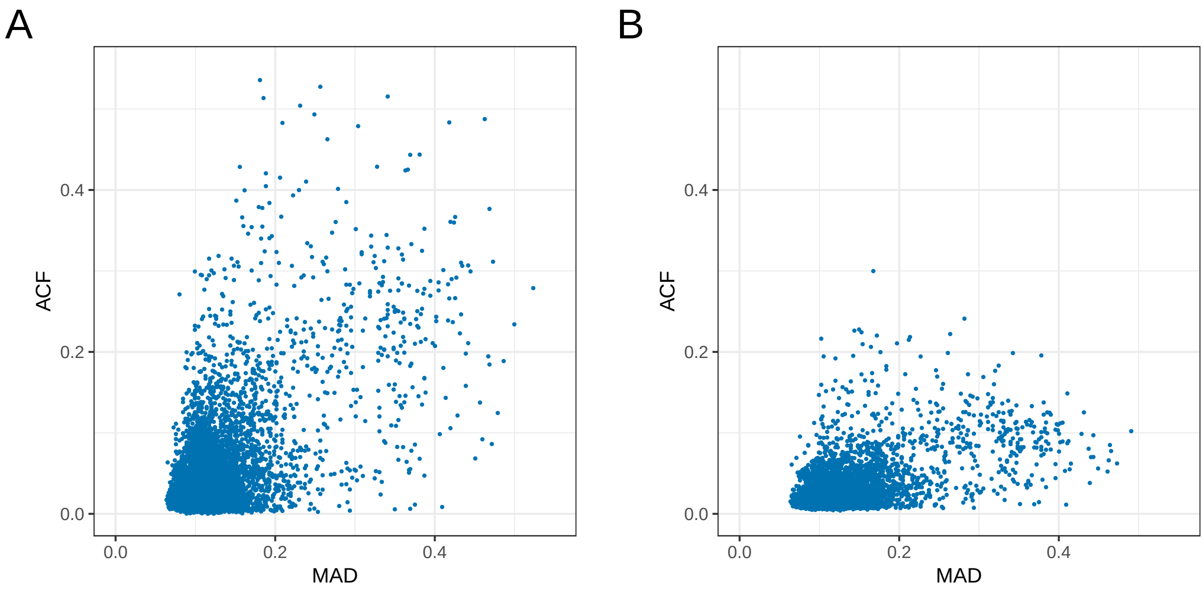


**Figure S1: Median absolute deviation and autocorrelation of autosomal $\log_2 R$ ratios.** After GC-correction, the majority of the samples have high data quality with median absolute deviations (MADs) less than 0.3 and lag-10 autocorrelations (ACFs) less than 0.1 (A). However, approximately 11% of the samples have autocorrelation greater than 0.1. To reduce autocorrelation, we applied a LOESS smoother to the scatterplot of genomic position versus $\log_2 R$ ratios using only SNPs that were likely heterozygous (0.4 $<$ BAF $<$ 0.6). From the LOESS model, we predicted the $\log_2 R$ ratios at all SNPs including those with BAFs outside the interval [0.4, 0.6]. The lag-10 autocorrelation of the residuals is less than 0.1 for nearly 98% of the PanC4 participants (B).

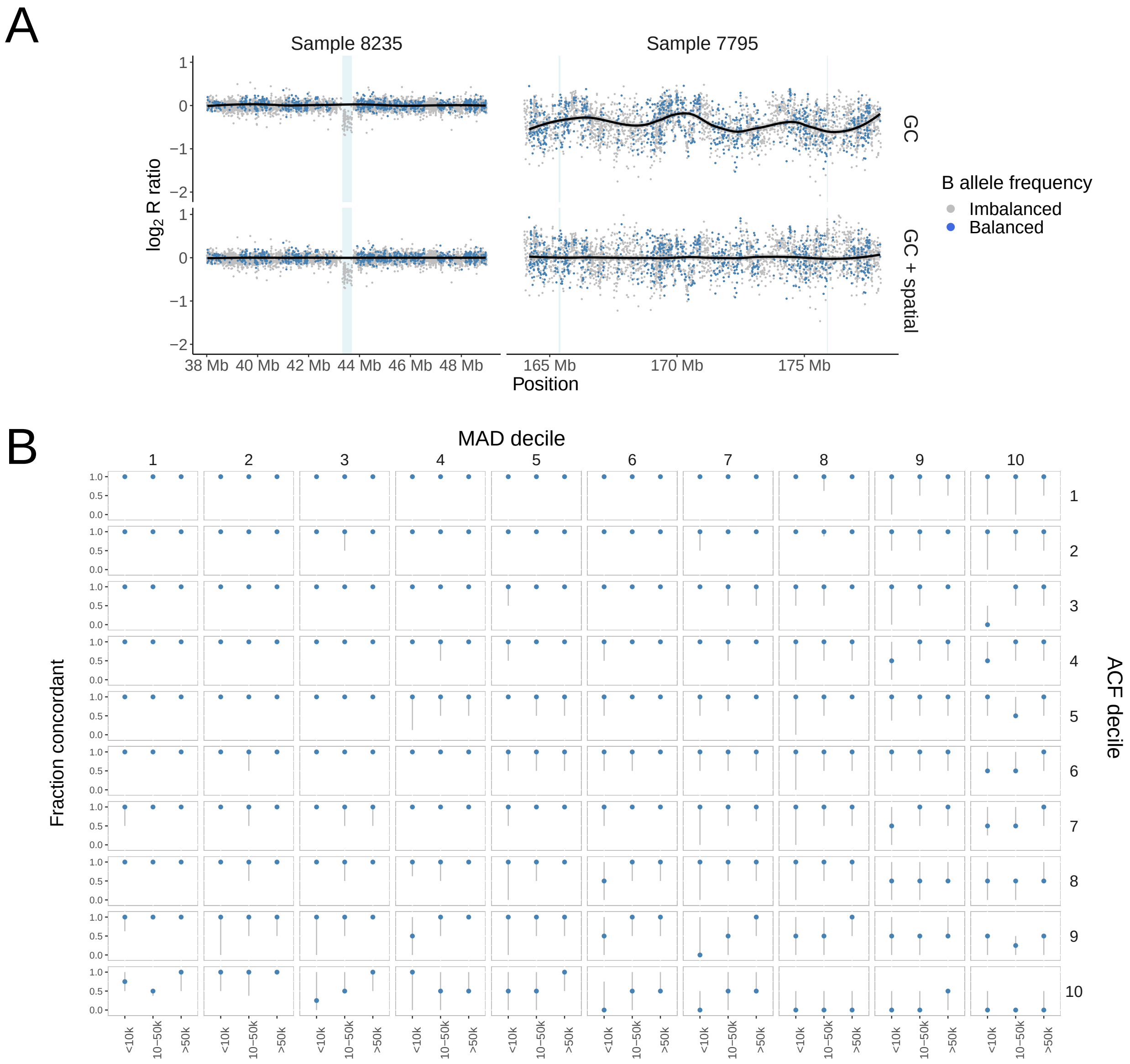


**Figure S2: Preprocessing and quality control analyses.** MAD and ACF measures of data quality among PanC4 participants were highly skewed to large values. **(A)** To reduce the ACF, we smoothed the $\log_2 R$ ratios with balanced B allele frequencies by genomic position using loess and predicted the $\log_2 R$ ratios in regions with imbalanced B allele frequencies from the loess model. The predicted $\log_2 R$ ratios for SNPs in the hemizygous deletion in Sample 8235 (44 Mb) are all near zero (solid black line), resulting in negative residuals in the GC+spatial panel (bottom). Shading indicates hemizygous deletions that were identified from a HMM fit to the GC-only and GC+spatially corrected data. **(B)** We summarized the median concordance (blue) and interquartile range (gray) of the CNV calls from the HMM before and after the spatial loess model for all 7,598 samples in strata of MAD and ACF decile. The additional preprocessing has a neglible effect on CNV inference in the high quality samples (top left), but substantial discordance is observed in the lower quality samples (bottom right) in part because of the larger number of false positives without the additional processing.

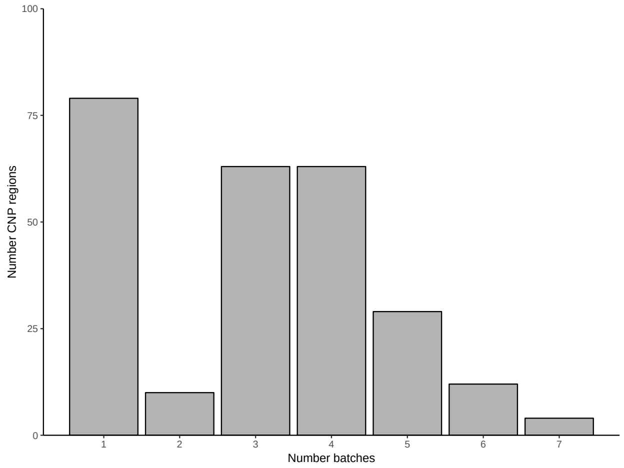


**Figure S3: Frequency of CNV regions with 1 to 7 batches identified by grouping the eCDFs of the $\log_2 R$ summaries.**

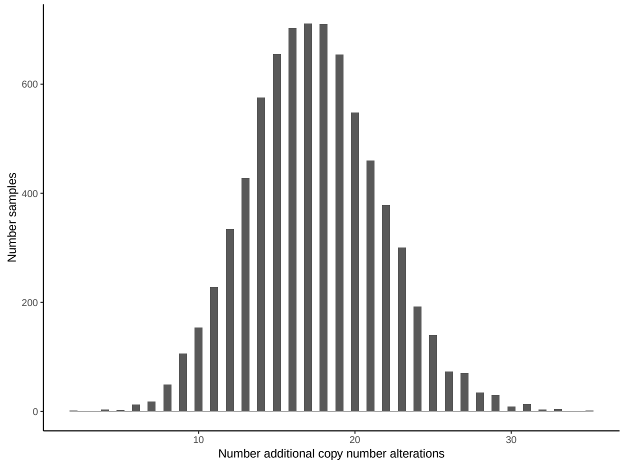


**Figure S4: Number of additional CNVs identified from the Bayesian mixture model.** On average, CNPBayes identified an additional 17 CNVs in each sample that were not detected from a hidden Markov model fit independently on the individual samples.

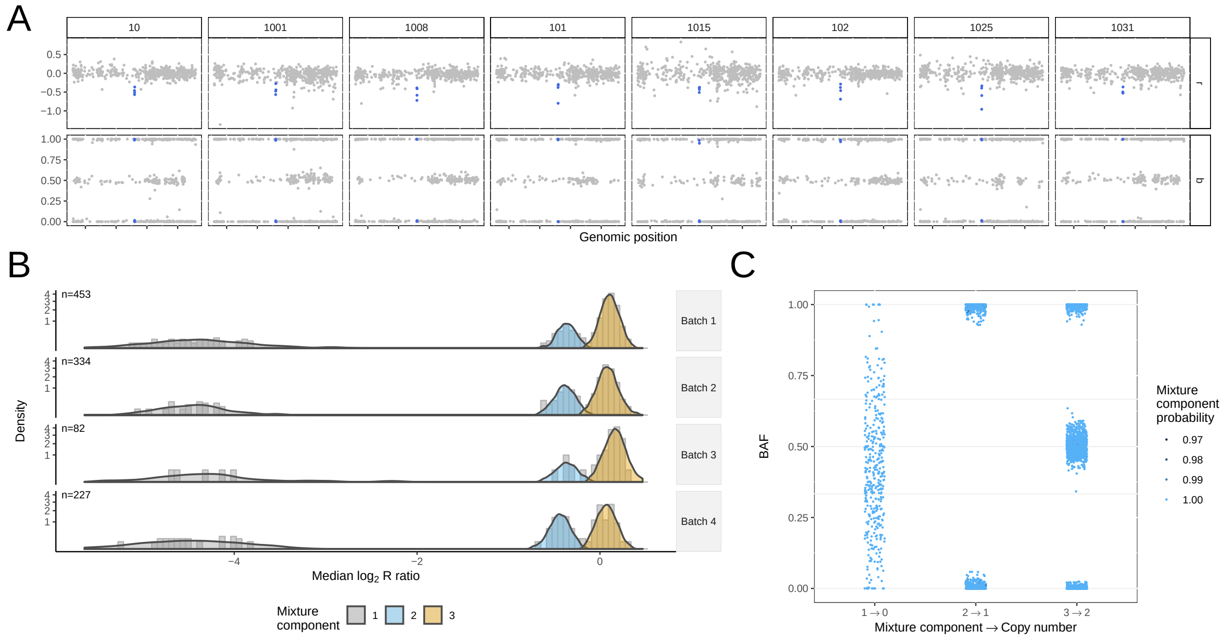


**Figure S5: Technical variation within and between samples obscures identification of hemizygous deletions. (A)** $\text{Log}_2$ R ratios (top) and B allele frequencies for SNPs in a 5kb CNP region (chr1:174,796,517-174,801,833) are highlighted in blue. The technical variation across the genome overwhelms the signal and the hemizygous deletion in these 10 samples is not detected. **(B)** The distribution of the median $\log_2 R$ for 6,038 samples. The lines of the histogram of the data is the density of the posterior predictive distribution with mixture components color-coded by the predicted copy number.

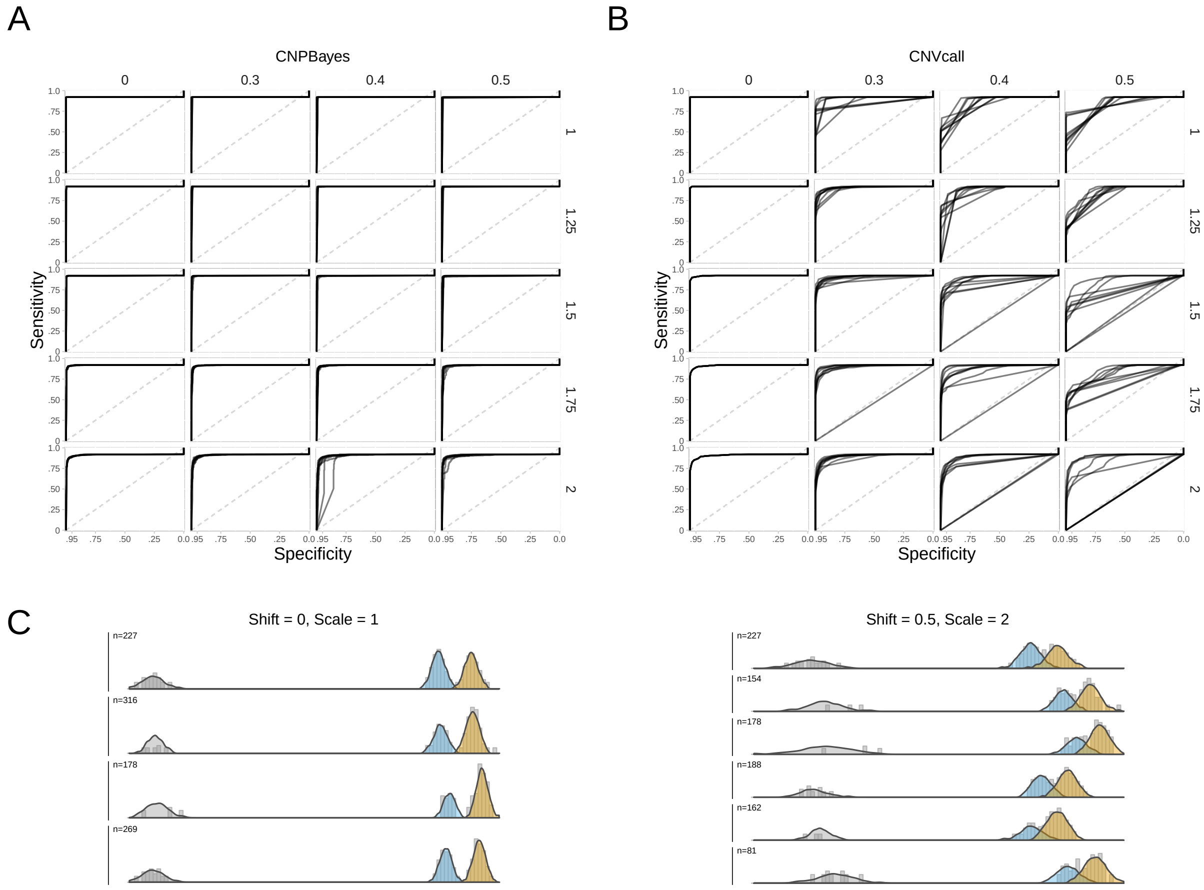


**Figure S6: Performance of CNV detection methods on HapMap data.** A deletion polymorphism for a 109 kb region on chromosome 4 in 990 HapMap samples that were processed on 16 chemistry plates. To simulate batch effects, a mean shift and/or rescaling of the variance was applied to the $\log_2 R$ ratios in a random subset of plates. **(A)** Each panel displays 10 receiver operator characteristic curves for CNPBayes evaluated on simulated datasets with the mean shift indicated in the column margins and scaling indicated in the row margins. While the true batch effect is not provided, `CNPBayes` attempts to infer the latent batches and has qualitatively similar performance as the level of difficulty increases from top-left to bottom-right. Two examples of the simulated data and posterior predictive distributions from `CNPBayes` are displayed in the top margin: no batch effect (left) and a mean shift of 0.5 with rescaling by factor of 2 (right). **(B)** By contrast, sensitivity and specificity of `CNVCALL` evaluated on the same data worsens as the simulated batch effects becomes more pronounced.
