## Supplemental Tables for "Bayesian copy number detection and association in large-scale studies"


PanC4\_workflow

- Home
- About
- License


### Supplemental Tables

###### Cristiano et al.

workflowr

- Summary
- Checks
- Past versions

**Last updated:** 2020-01-17

**Checks:**  5  2

**Knit directory:** `PanC4_workflow/`

This reproducible R Markdown analysis was created with workflowr (version 1.6.0). The *Checks* tab describes the reproducibility checks that were applied when the results were created. The *Past versions* tab lists the development history.

---

**R Markdown file:** uncommitted changes

The R Markdown is untracked by Git. To know which version of the R Markdown file created these results, you’ll want to first commit it to the Git repo. If you’re still working on the analysis, you can ignore this warning. When you’re finished, you can run `wflow_publish` to commit the R Markdown file and build the HTML.

**Environment:** objects present

The global environment had objects present when the code in the R Markdown file was run. These objects can affect the analysis in your R Markdown file in unknown ways. For reproduciblity it’s best to always run the code in an empty environment. Use `wflow_publish` or `wflow_build` to ensure that the code is always run in an empty environment.

The following objects were defined in the global environment when these results were created:

| Name | Class | Size |
| --- | --- | --- |
| rv | list | 2.9 Kb |

**Seed:** `set.seed(20190721)`

The command `set.seed(20190721)` was run prior to running the code in the R Markdown file. Setting a seed ensures that any results that rely on randomness, e.g. subsampling or permutations, are reproducible.

**Session information:** recorded

Great job! Recording the operating system, R version, and package versions is critical for reproducibility.

**Cache:** none

Nice! There were no cached chunks for this analysis, so you can be confident that you successfully produced the results during this run.

**File paths:** relative

Great job! Using relative paths to the files within your workflowr project makes it easier to run your code on other machines.

**Repository version:** 0679044

Great! You are using Git for version control. Tracking code development and connecting the code version to the results is critical for reproducibility. The version displayed above was the version of the Git repository at the time these results were generated.   
  
 Note that you need to be careful to ensure that all relevant files for the analysis have been committed to Git prior to generating the results (you can use `wflow_publish` or `wflow_git_commit`). workflowr only checks the R Markdown file, but you know if there are other scripts or data files that it depends on. Below is the status of the Git repository when the results were generated:

```
Ignored files:
    Ignored:    .Rhistory
    Ignored:    analysis/.Rhistory
    Ignored:    analysis/gwa_cache/
    Ignored:    analysis/overview_v2.html
    Ignored:    code/.Rhistory
    Ignored:    output/batch_effects.Rmd/
    Ignored:    output/convergence_diagnostics_nobatch.Rmd/
    Ignored:    output/convergence_noz.Rmd/

Untracked files:
    Untracked:  analysis/Rplots.pdf
    Untracked:  analysis/approach.Rmd
    Untracked:  analysis/archive/
    Untracked:  analysis/association_figures.Rmd
    Untracked:  analysis/batch_effects.Rmd
    Untracked:  analysis/cnp_regions.Rmd
    Untracked:  analysis/cnv_effect_size.Rmd
    Untracked:  analysis/cnv_vaf.Rmd
    Untracked:  analysis/copynumber.Rmd
    Untracked:  analysis/gwa.Rmd
    Untracked:  analysis/overview_v2.Rmd
    Untracked:  code/bayesfactors_old.Rmd
    Untracked:  code/cn_comparison.Rmd
    Untracked:  code/convergence_diagnostics.Rmd
    Untracked:  code/convergence_diagnostics.html
    Untracked:  code/convergence_diagnostics_interaction.Rmd
    Untracked:  code/convergence_diagnostics_interaction.html
    Untracked:  code/convergence_diagnostics_nobatch.Rmd
    Untracked:  code/convergence_noz.Rmd
    Untracked:  code/effective_size.Rmd
    Untracked:  code/figure/
    Untracked:  code/old/
    Untracked:  code/output/
    Untracked:  code/panc.funs/
    Untracked:  code/panc.funs_0.0.2.tar.gz
    Untracked:  code/scratch.Rmd
    Untracked:  code/snp_assoc.Rmd
    Untracked:  data/cnv_assoc_interaction.R/
    Untracked:  data/collect_gwa.R/
    Untracked:  data/snp_association4.R/
    Untracked:  output/approach.Rmd/
    Untracked:  output/association_figures.Rmd/
    Untracked:  output/cnv_effect_size.Rmd/
    Untracked:  output/cnv_vaf.Rmd/
    Untracked:  output/collect_cnv_association.R/
    Untracked:  output/collect_gwa.R/
    Untracked:  output/convergence_diagnostics.Rmd/
    Untracked:  output/convergence_diagnostics_interaction.Rmd/
    Untracked:  output/copynumber.Rmd/
    Untracked:  output/data/
    Untracked:  output/figures/
    Untracked:  output/gwa.Rmd/
    Untracked:  output/overview.Rmd/
    Untracked:  output/overview_v2.Rmd/
    Untracked:  output/snp_assoc.Rmd/
    Untracked:  tmp.pdf

Unstaged changes:
    Modified:   .Rprofile
    Modified:   README.md
    Modified:   _workflowr.yml
    Modified:   analysis/_site.yml
    Modified:   analysis/about.Rmd
    Modified:   analysis/index.Rmd
    Modified:   analysis/license.Rmd
    Modified:   code/README.md
    Modified:   data/README.md
    Modified:   output/README.md
```

Note that any generated files, e.g. HTML, png, CSS, etc., are not included in this status report because it is ok for generated content to have uncommitted changes.

---

There are no past versions. Publish this analysis with `wflow_publish()` to start tracking its development.

---

### Table S1: Copy number variant (CNV) regions

Genomic regions from UCSC genome build hg18 evaluated by CNPBayes as possible copy number polymorphisms.

| Region id | Chr | Start | End | Width | Number probes | 1000G genotypes | European allele frequency | 1000G variant id | Reference |
| --- | --- | --- | --- | --- | --- | --- | --- | --- | --- |
| CNP\_001 | chr1 | 1,627,805 | 1,673,809 | 46,005 | 8 | CN0, CN2 | 0.044, 0.024 | esv3585014, esv3585015 | 1000G |
| CNP\_002 | chr1 | 12,854,895 | 12,919,194 | 64,300 | 19 | CN0 | 0.029 | esv3585241 | 1000G |
| CNP\_003 | chr1 | 12,856,327 | 12,893,193 | 36,867 | 10 | CN0, CN2 | 0.033, 0.008 | esv3585242, esv3585243 | 1000G |
| CNP\_004 | chr1 | 12,877,893 | 12,933,272 | 55,380 | 31 | CN0 | 0.025 | esv3585244 | 1000G |
| CNP\_005 | chr1 | 12,893,536 | 12,948,772 | 55,237 | 28 | CN0 | 0.032 | esv3585245 | 1000G |
| CNP\_007 | chr1 | 12,901,370 | 12,921,250 | 19,881 | 22 | CN0, CN2, CN3 | 0.039, 0.039, 0.001 | esv3585247, esv3585248, esv3585249 | 1000G |
| CNP\_010 | chr1 | 17,238,417 | 17,266,696 | 28,280 | 18 | CN2, CN3 | 0.462, 0.007 | esv3585353, esv3585354 | 1000G |
| CNP\_011 | chr1 | 17,263,114 | 17,273,513 | 10,400 | 7 | CN2, CN3 | 0.463, 0.006 | esv3585355, esv3585356 | 1000G |
| CNP\_012 | chr1 | 22,307,650 | 22,332,441 | 24,792 | 9 | CN0 | 0.024 | esv3585459 | 1000G |
| CNP\_013 | chr1 | 22,316,148 | 22,334,511 | 18,364 | 4 | CN0, CN2 | 0.027, 0.001 | esv3585460, esv3585461 | 1000G |
| CNP\_014 | chr1 | 25,592,642 | 25,661,222 | 68,581 | 9 | CN0, CN2 | 0.4, 0.002 | esv3585521, esv3585522 | 1000G |
| CNP\_015 | chr1 | 28,206,060 | 28,261,986 | 55,927 | 37 | CN0, CN2 | 0.002, 0 | esv3585564, esv3585565 | 1000G |
| CNP\_016 | chr1 | 104,102,743 | 104,136,676 | 33,934 | 18 | CN2, CN3 | 0.074, 0.001 | esv3587012, esv3587013 | 1000G |
| CNP\_018 | chr1 | 109,366,972 | 109,372,000 | 5,029 | 6 | CN0 | 0.025 | esv3587138 | 1000G |
| CNP\_019 | chr1 | 110,225,019 | 110,245,280 | 20,262 | 8 | CN0 | 0.723 | esv3587154 | 1000G |
| CNP\_021 | chr1 | 151,333,216 | 151,413,627 | 80,412 | 50 | CN0, CN2 | 0.006, 0 | esv3587526, esv3587527 | 1000G |
| CNP\_022 | chr1 | 174,796,517 | 174,801,833 | 5,317 | 4 | CN0 | 0.113 | esv3587968 | 1000G |
| CNP\_023 | chr1 | 196,801,025 | 196,892,322 | 91,298 | 27 | NA | NA |  | Panc4 |
| CNP\_024 | chr1 | 240,137,922 | 240,302,859 | 164,938 | 71 | CN2 | 0.002 | esv3589293 | 1000G |
| CNP\_025 | chr1 | 248,756,741 | 248,797,597 | 40,857 | 6 | CN0, CN2 | 0.124, 0.001 | esv3589559, esv3589560 | 1000G |
| CNP\_026 | chr2 | 41,238,373 | 41,250,303 | 11,931 | 5 | CN0 | 0.066 | esv3590473 | 1000G |
| CNP\_027 | chr2 | 49,532,573 | 49,542,423 | 9,851 | 4 | CN0 | 0.045 | esv3590687 | 1000G |
| CNP\_028 | chr2 | 51,926,595 | 51,927,817 | 1,223 | 5 | CN0 | 0.117 | esv3590770 | 1000G |
| CNP\_030 | chr2 | 90,066,423 | 90,120,854 | 54,432 | 6 | CN0, CN2 | 0.032, 0.005 | esv3591623, esv3591624 | 1000G |
| CNP\_031 | chr2 | 90,212,408 | 90,249,248 | 36,841 | 4 | CN0, CN2 | 0.039, 0.005 | esv3591627, esv3591628 | 1000G |
| CNP\_032 | chr2 | 130,549,320 | 130,558,827 | 9,508 | 4 | CN0 | 0 | esv3592431 | 1000G |
| CNP\_033 | chr2 | 195,979,320 | 195,984,295 | 4,976 | 4 | CN0 | 0.063 | esv3593856 | 1000G |
| CNP\_034 | chr2 | 208,351,101 | 208,359,313 | 8,213 | 4 | CN0 | 0.163 | esv3594087 | 1000G |
| CNP\_035 | chr2 | 213,183,956 | 213,192,007 | 8,052 | 8 | CN0 | 0.062 | esv3594201 | 1000G |
| CNP\_036 | chr2 | 242,917,734 | 243,044,147 | 126,414 | 22 | NA | NA |  | Panc4 |
| CNP\_037 | chr3 | 37,978,345 | 37,986,932 | 8,588 | 6 | CN0 | 0.088 | esv3595902 | 1000G |
| CNP\_038 | chr3 | 65,188,826 | 65,214,723 | 25,898 | 11 | CN0 | 0.078 | esv3596416 | 1000G |
| CNP\_040 | chr3 | 75,428,675 | 75,703,846 | 275,172 | 12 | NA | NA |  | Panc4 |
| CNP\_041 | chr3 | 148,963,333 | 148,969,710 | 6,378 | 6 | CN0 | 0.036 | esv3598142 | 1000G |
| CNP\_042 | chr3 | 162,127,885 | 162,143,633 | 15,749 | 7 | CN0 | 0.071 | esv3598432 | 1000G |
| CNP\_043 | chr3 | 162,213,371 | 162,235,756 | 22,386 | 4 | CN0 | 0.003 | esv3598433 | 1000G |
| CNP\_044 | chr3 | 173,239,453 | 173,289,281 | 49,829 | 15 | NA | NA |  | Panc4 |
| CNP\_045 | chr3 | 191,064,690 | 191,071,659 | 6,970 | 5 | CN0 | 0.258 | esv3599104 | 1000G |
| CNP\_046 | chr3 | 191,988,492 | 191,997,177 | 8,686 | 6 | CN2 | 0.002 | esv3599142 | 1000G |
| CNP\_047 | chr3 | 193,135,464 | 193,142,938 | 7,475 | 4 | CN0 | 0.027 | esv3599184 | 1000G |
| CNP\_048 | chr3 | 195,452,012 | 195,457,240 | 5,229 | 4 | CN2, CN3, CN4 | 0.444, 0.002, 0 | esv3599243, esv3599244, esv3599245 | 1000G |
| CNP\_049 | chr3 | 195,954,431 | 196,022,808 | 68,378 | 33 | CN0, CN2 | 0.011, 0.001 | esv3599276, esv3599277 | 1000G |
| CNP\_050 | chr4 | 3,446,697 | 3,451,946 | 5,250 | 16 | CN0, CN2 | 0.003, 0.028 | esv3599429, esv3599430 | 1000G |
| CNP\_051 | chr4 | 9,370,866 | 9,410,140 | 39,275 | 4 | CN0, CN2, CN3 | 0.002, 0.007, 0.001 | esv3599568, esv3599569, esv3599570 | 1000G |
| CNP\_052 | chr4 | 9,418,201 | 9,457,405 | 39,205 | 5 | CN0, CN2, CN3 | 0.002, 0.006, 0.001 | esv3599572, esv3599573, esv3599574 | 1000G |
| CNP\_053 | chr4 | 10,392,397 | 10,402,099 | 9,703 | 4 | CN0, CN2 | 0.252, 0 | esv3599620, esv3599621 | 1000G |
| CNP\_054 | chr4 | 34,781,317 | 34,849,886 | 68,570 | 6 | CN0 | 0.297 | esv3600181 | 1000G |
| CNP\_055 | chr4 | 64,133,825 | 64,153,198 | 19,374 | 5 | CN0, CN2 | 0.092, 0 | esv3600733, esv3600734 | 1000G |
| CNP\_056 | chr4 | 69,375,591 | 69,491,543 | 115,953 | 7 | CN0 | 0.368 | esv3600874 | 1000G |
| CNP\_057 | chr4 | 70,122,981 | 70,231,746 | 108,766 | 4 | CN0 | 0.154 | esv3600896 | 1000G |
| CNP\_059 | chr4 | 122,282,235 | 122,289,973 | 7,739 | 5 | CN0 | 0.055 | esv3602006 | 1000G |
| CNP\_060 | chr4 | 151,876,054 | 151,885,557 | 9,504 | 4 | CN0, CN2 | 0.001, 0.001 | esv3602668, esv3602669 | 1000G |
| CNP\_061 | chr4 | 152,790,167 | 152,794,773 | 4,607 | 5 | CN0 | 0.056 | esv3602694 | 1000G |
| CNP\_062 | chr4 | 161,059,762 | 161,071,059 | 11,298 | 13 | NA | NA |  | Panc4 |
| CNP\_064 | chr5 | 814,446 | 825,367 | 10,922 | 14 | CN0, CN2, CN3, CN4 | 0.04, 0.046, 0.003, 0.002 | esv3603782, esv3603783, esv3603784, esv3603785 | 1000G |
| CNP\_065 | chr5 | 32,107,381 | 32,159,517 | 52,137 | 32 | NA | NA |  | Panc4 |
| CNP\_066 | chr5 | 45,983,680 | 49,522,346 | 3,538,667 | 27 | NA | NA |  | Panc4 |
| CNP\_067 | chr5 | 57,320,371 | 57,338,133 | 17,763 | 5 | CN0 | 0.777 | esv3605146 | 1000G |
| CNP\_069 | chr5 | 70,306,516 | 70,313,302 | 6,787 | 9 | CN0, CN2, CN3 | 0.028, 0.086, 0.015 | esv3605386, esv3605387, esv3605388 | 1000G |
| CNP\_070 | chr5 | 97,048,466 | 97,099,320 | 50,855 | 18 | NA | NA |  | Panc4 |
| CNP\_071 | chr5 | 110,903,828 | 110,913,774 | 9,947 | 4 | CN0 | 0 | esv3606276 | 1000G |
| CNP\_072 | chr5 | 140,222,274 | 140,239,347 | 17,074 | 29 | CN0 | 0.076 | esv3606958 | 1000G |
| CNP\_074 | chr5 | 140,554,408 | 140,558,942 | 4,535 | 10 | CN0, CN2, CN3, CN4, CN5, CN6, CN7 | 0.015, 0.094, 0.307, 0.015, 0.005, 0.002, 0.007 | esv3606964, esv3606965, esv3606966, esv3606967, esv3606968, esv3606969, esv3606970 | 1000G |
| CNP\_075 | chr5 | 142,174,919 | 142,260,351 | 85,433 | 33 | CN0, CN2 | 0.009, 0 | esv3607010, esv3607011 | 1000G |
| CNP\_076 | chr5 | 142,263,109 | 142,447,062 | 183,954 | 55 | CN0, CN2 | 0.002, 0 | esv3607012, esv3607013 | 1000G |
| CNP\_077 | chr5 | 151,511,011 | 151,518,882 | 7,872 | 6 | CN0 | 0.141 | esv3607195 | 1000G |
| CNP\_078 | chr5 | 151,514,809 | 151,518,869 | 4,061 | 4 | CN0 | 0.164 | esv3607196 | 1000G |
| CNP\_079 | chr5 | 155,475,886 | 155,488,649 | 12,764 | 4 | CN0, CN2 | 0.198, 0.002 | esv3607277, esv3607278 | 1000G |
| CNP\_080 | chr6 | 327,586 | 368,712 | 41,127 | 11 | CN2, CN3, CN4 | 0.797, 0.022, 0.004 | esv3607848, esv3607849, esv3607850 | 1000G |
| CNP\_081 | chr6 | 29,851,172 | 29,904,515 | 53,344 | 24 | CN0 | 0.111 | esv3608493 | 1000G |
| CNP\_083 | chr6 | 30,994,015 | 30,995,083 | 1,069 | 12 | CN0 | 0.145 | esv3608521 | 1000G |
| CNP\_084 | chr6 | 31,131,451 | 31,272,307 | 140,857 | 363 | CN0, CN2 | 0.002, 0 | esv3608531, esv3608532 | 1000G |
| CNP\_085 | chr6 | 31,219,515 | 31,229,793 | 10,279 | 8 | CN0 | 0.023 | esv3608543 | 1000G |
| CNP\_086 | chr6 | 31,275,976 | 31,279,146 | 3,171 | 9 | CN0 | 0.101 | esv3608549 | 1000G |
| CNP\_087 | chr6 | 31,337,847 | 31,341,985 | 4,139 | 12 | CN0 | 0.013 | esv3608560 | 1000G |
| CNP\_088 | chr6 | 32,604,936 | 32,616,132 | 11,197 | 13 | CN0, CN2 | 0.406, 0.317 | esv3608602, esv3608603 | 1000G |
| CNP\_089 | chr6 | 35,521,984 | 35,568,895 | 46,912 | 14 | CN0, CN2 | 0.002, 0.001 | esv3608684, esv3608685 | 1000G |
| CNP\_090 | chr6 | 35,754,684 | 35,766,728 | 12,045 | 9 | CN2, CN3 | 0.491, 0.019 | esv3608692, esv3608693 | 1000G |
| CNP\_091 | chr6 | 67,009,228 | 67,049,033 | 39,806 | 10 | CN0 | 0.076 | esv3609317 | 1000G |
| CNP\_093 | chr6 | 78,879,305 | 78,898,720 | 19,416 | 4 | CN0, CN2 | 0, 0 | esv3609655, esv3609656 | 1000G |
| CNP\_094 | chr6 | 78,892,808 | 79,053,430 | 160,623 | 37 | CN0, CN2 | 0.267, 0 | esv3609657, esv3609658 | 1000G |
| CNP\_095 | chr6 | 78,967,097 | 79,036,275 | 69,179 | 12 | CN0, CN2 | 0.265, 0 | esv3609663, esv3609664 | 1000G |
| CNP\_097 | chr6 | 81,283,720 | 81,293,577 | 9,858 | 4 | CN0 | 0.055 | esv3609714 | 1000G |
| CNP\_098 | chr6 | 93,575,094 | 93,579,998 | 4,905 | 6 | CN0 | 0 | esv3609976 | 1000G |
| CNP\_099 | chr6 | 168,364,813 | 168,593,956 | 229,144 | 172 | NA | NA |  | Panc4 |
| CNP\_100 | chr7 | 39,119 | 75,937 | 36,819 | 13 | CN2 | 0.039 | esv3611744 | 1000G |
| CNP\_102 | chr7 | 7,048,022 | 7,053,955 | 5,934 | 4 | CN0 | 0.004 | esv3612059 | 1000G |
| CNP\_103 | chr7 | 61,994,170 | 62,397,642 | 403,473 | 21 | NA | NA |  | Panc4 |
| CNP\_104 | chr7 | 62,687,405 | 62,723,597 | 36,193 | 4 | CN0, CN2, CN3 | 0.002, 0.009, 0.001 | esv3613391, esv3613392, esv3613393 | 1000G |
| CNP\_105 | chr7 | 76,417,751 | 76,615,349 | 197,599 | 21 | NA | NA |  | Panc4 |
| CNP\_106 | chr7 | 91,031,079 | 91,042,590 | 11,512 | 4 | CN0 | 0 | esv3614118 | 1000G |
| CNP\_107 | chr7 | 100,675,367 | 100,685,330 | 9,964 | 83 | NA | NA |  | Panc4 |
| CNP\_108 | chr7 | 119,153,885 | 119,162,547 | 8,663 | 5 | CN0 | 0.001 | esv3614766 | 1000G |
| CNP\_109 | chr7 | 141,765,307 | 141,792,849 | 27,543 | 5 | CN0, CN2 | 0.197, 0.01 | esv3615267, esv3615268 | 1000G |
| CNP\_110 | chr7 | 142,475,484 | 142,486,103 | 10,620 | 4 | CN0 | 0.412 | esv3615291 | 1000G |
| CNP\_111 | chr7 | 142,827,954 | 142,881,540 | 53,587 | 35 | NA | NA |  | Panc4 |
| CNP\_112 | chr8 | 2,129,510 | 2,135,062 | 5,553 | 5 | CN0 | 0.001 | esv3615861 | 1000G |
| CNP\_113 | chr8 | 3,786,311 | 3,790,617 | 4,307 | 4 | CN0, CN2 | 0.055, 0 | esv3615928, esv3615929 | 1000G |
| CNP\_114 | chr8 | 3,996,836 | 4,004,485 | 7,650 | 6 | CN0, CN2 | 0.002, 0.001 | esv3615947, esv3615948 | 1000G |
| CNP\_115 | chr8 | 5,595,438 | 5,605,648 | 10,211 | 10 | CN0, CN2 | 0.084, 0 | esv3616046, esv3616047 | 1000G |
| CNP\_117 | chr8 | 13,613,298 | 13,650,857 | 37,560 | 15 | CN0, CN2 | 0, 0.001 | esv3616334, esv3616335 | 1000G |
| CNP\_118 | chr8 | 13,625,663 | 13,640,463 | 14,801 | 6 | CN0 | 0 | esv3616336 | 1000G |
| CNP\_119 | chr8 | 13,626,465 | 13,652,740 | 26,276 | 7 | CN0 | 0 | esv3616337 | 1000G |
| CNP\_120 | chr8 | 13,635,690 | 13,645,420 | 9,731 | 4 | CN0 | 0 | esv3616338 | 1000G |
| CNP\_121 | chr8 | 15,401,664 | 15,414,791 | 13,128 | 7 | CN0 | 0.025 | esv3616406 | 1000G |
| CNP\_122 | chr8 | 16,262,082 | 16,274,699 | 12,618 | 5 | CN0, CN2 | 0.037, 0 | esv3616450, esv3616451 | 1000G |
| CNP\_123 | chr8 | 39,195,825 | 39,389,230 | 193,406 | 5 | CN0 | 0.428 | esv3616947 | 1000G |
| CNP\_124 | chr8 | 40,182,839 | 40,189,804 | 6,966 | 6 | CN0 | 0.056 | esv3616969 | 1000G |
| CNP\_125 | chr8 | 85,260,961 | 85,269,169 | 8,209 | 4 | CN0 | 0.037 | esv3617854 | 1000G |
| CNP\_126 | chr8 | 120,019,604 | 120,027,755 | 8,152 | 4 | CN0 | 0 | esv3618585 | 1000G |
| CNP\_127 | chr8 | 135,059,548 | 135,068,056 | 8,509 | 6 | CN0 | 0.001 | esv3618901 | 1000G |
| CNP\_128 | chr8 | 137,682,484 | 137,857,327 | 174,844 | 50 | NA | NA |  | Panc4 |
| CNP\_129 | chr9 | 15,157,294 | 15,159,553 | 2,260 | 5 | CN0 | 0 | esv3619764 | 1000G |
| CNP\_130 | chr9 | 72,030,971 | 72,054,480 | 23,510 | 5 | CN2 | 0.005 | esv3620559 | 1000G |
| CNP\_131 | chr9 | 132,463,983 | 132,648,102 | 184,120 | 115 | CN0, CN2 | 0.004, 0 | esv3621839, esv3621840 | 1000G |
| CNP\_132 | chr9 | 132,654,988 | 132,798,900 | 143,913 | 30 | CN0, CN2 | 0.016, 0 | esv3621845, esv3621846 | 1000G |
| CNP\_133 | chr9 | 134,293,849 | 134,452,956 | 159,108 | 127 | CN0, CN2 | 0.018, 0 | esv3621887, esv3621888 | 1000G |
| CNP\_134 | chr9 | 135,938,579 | 135,956,026 | 17,448 | 13 | CN0, CN2 | 0.001, 0.042 | esv3621931, esv3621932 | 1000G |
| CNP\_135 | chr10 | 20,849,543 | 20,860,166 | 10,624 | 5 | CN0 | 0.237 | esv3622561 | 1000G |
| CNP\_136 | chr10 | 47,531,022 | 47,599,037 | 68,016 | 10 | CN0, CN2, CN3 | 0.009, 0.041, 0.003 | esv3623125, esv3623126, esv3623127 | 1000G |
| CNP\_138 | chr10 | 47,596,038 | 47,634,490 | 38,453 | 20 | CN0, CN2, CN3 | 0.009, 0.041, 0.003 | esv3623128, esv3623129, esv3623130 | 1000G |
| CNP\_139 | chr10 | 47,645,964 | 47,697,149 | 51,186 | 18 | CN0, CN2, CN3 | 0.002, 0.044, 0.001 | esv3623133, esv3623134, esv3623135 | 1000G |
| CNP\_140 | chr10 | 58,512,456 | 58,527,377 | 14,922 | 4 | CN0 | 0.051 | esv3623398 | 1000G |
| CNP\_141 | chr10 | 68,078,481 | 68,114,481 | 36,001 | 18 | NA | NA |  | Panc4 |
| CNP\_142 | chr10 | 90,248,794 | 90,448,452 | 199,659 | 67 | CN0, CN2 | 0, 0 | esv3624131, esv3624132 | 1000G |
| CNP\_143 | chr10 | 90,551,092 | 90,632,203 | 81,112 | 49 | CN0, CN2 | 0.001, 0 | esv3624140, esv3624141 | 1000G |
| CNP\_144 | chr10 | 122,769,703 | 122,787,503 | 17,801 | 9 | CN0 | 0 | esv3624746 | 1000G |
| CNP\_145 | chr10 | 124,339,615 | 124,377,666 | 38,052 | 7 | CN0 | 0.001 | esv3624776 | 1000G |
| CNP\_147 | chr10 | 135,348,035 | 135,378,260 | 30,226 | 69 | CN0, CN2, CN3 | 0, 0.014, 0 | esv3625052, esv3625053, esv3625054 | 1000G |
| CNP\_148 | chr11 | 1,015,786 | 1,019,413 | 3,628 | 67 | NA | NA |  | Panc4 |
| CNP\_149 | chr11 | 3,238,736 | 3,244,087 | 5,352 | 6 | CN0 | 0.037 | esv3625135 | 1000G |
| CNP\_150 | chr11 | 3,238,750 | 3,361,018 | 122,269 | 38 | CN2, CN3 | 0.037, 0.001 | esv3625136, esv3625137 | 1000G |
| CNP\_151 | chr11 | 5,873,986 | 5,883,494 | 9,509 | 8 | CN0 | 0.001 | esv3625252 | 1000G |
| CNP\_152 | chr11 | 7,812,280 | 7,832,579 | 20,300 | 7 | CN0, CN2 | 0.034, 0 | esv3625301, esv3625302 | 1000G |
| CNP\_153 | chr11 | 18,941,736 | 18,963,997 | 22,262 | 6 | CN0, CN2, CN3, CN4 | 0.044, 0.193, 0.031, 0.021 | esv3625519, esv3625520, esv3625521, esv3625522 | 1000G |
| CNP\_154 | chr11 | 42,967,415 | 42,976,408 | 8,994 | 5 | CN0, CN2, CN3, CN4 | 0.005, 0.169, 0, 0.001 | esv3626131, esv3626132, esv3626133, esv3626134 | 1000G |
| CNP\_155 | chr11 | 55,368,372 | 55,426,969 | 58,598 | 13 | CN0, CN2 | 0.284, 0.001 | esv3626445, esv3626446 | 1000G |
| CNP\_157 | chr11 | 58,814,995 | 58,855,468 | 40,474 | 5 | CN0, CN2 | 0, 0.005 | esv3626529, esv3626530 | 1000G |
| CNP\_158 | chr11 | 81,500,537 | 81,521,332 | 20,796 | 9 | CN0 | 0.097 | esv3626968 | 1000G |
| CNP\_159 | chr11 | 86,304,229 | 86,306,559 | 2,331 | 6 | CN0 | 0.092 | esv3627094 | 1000G |
| CNP\_160 | chr11 | 132,953,768 | 133,158,580 | 204,813 | 85 | CN2 | 0.009 | esv3628139 | 1000G |
| CNP\_161 | chr11 | 134,621,447 | 134,725,032 | 103,586 | 50 | CN2 | 0.01 | esv3628188 | 1000G |
| CNP\_162 | chr12 | 886,071 | 967,457 | 81,387 | 12 | CN0, CN2 | 0.001, 0 | esv3628256, esv3628257 | 1000G |
| CNP\_163 | chr12 | 8,000,912 | 8,114,429 | 113,518 | 40 | NA | NA |  | Panc4 |
| CNP\_164 | chr12 | 8,360,756 | 8,389,845 | 29,090 | 4 | CN2, CN3 | 0.149, 0.02 | esv3628458, esv3628459 | 1000G |
| CNP\_165 | chr12 | 9,915,398 | 10,038,593 | 123,196 | 50 | CN0, CN2 | 0.005, 0 | DUP\_uwash\_chr12\_9915398\_10038593 | 1000G |
| CNP\_166 | chr12 | 11,222,191 | 11,249,671 | 27,481 | 6 | CN0, CN2 | 0.491, 0 | esv3628561, esv3628562 | 1000G |
| CNP\_167 | chr12 | 27,648,174 | 27,655,202 | 7,029 | 7 | CN0, CN2 | 0.026, 0 | esv3628941, esv3628942 | 1000G |
| CNP\_168 | chr12 | 31,266,287 | 31,407,303 | 141,017 | 37 | NA | NA |  | Panc4 |
| CNP\_169 | chr12 | 33,296,520 | 33,307,372 | 10,853 | 4 | CN0 | 0.221 | esv3629097 | 1000G |
| CNP\_170 | chr12 | 37,998,057 | 38,371,887 | 373,831 | 21 | NA | NA |  | Panc4 |
| CNP\_171 | chr12 | 53,086,320 | 53,087,693 | 1,374 | 6 | CN0 | 0.08 | esv3629525 | 1000G |
| CNP\_172 | chr12 | 59,935,796 | 59,952,863 | 17,068 | 4 | CN0 | 0.17 | esv3629660 | 1000G |
| CNP\_173 | chr12 | 70,679,288 | 70,682,287 | 3,000 | 4 | CN0 | 0.028 | esv3629885 | 1000G |
| CNP\_174 | chr12 | 70,872,239 | 70,878,209 | 5,971 | 4 | CN0 | 0.164 | esv3629893 | 1000G |
| CNP\_175 | chr12 | 80,153,371 | 80,162,648 | 9,278 | 4 | CN0, CN2, CN3 | 0, 0.261, 0.011 | esv3630087, esv3630088, esv3630089 | 1000G |
| CNP\_176 | chr12 | 121,500,914 | 121,595,375 | 94,462 | 31 | CN0, CN2 | 0.005, 0 | esv3630934, esv3630935 | 1000G |
| CNP\_177 | chr12 | 124,495,947 | 124,500,196 | 4,250 | 5 | CN2 | 0.1 | esv3631000 | 1000G |
| CNP\_178 | chr12 | 127,430,503 | 127,812,976 | 382,474 | 141 | CN0, CN2 | 0, 0.008 | esv3631094, esv3631095 | 1000G |
| CNP\_179 | chr12 | 127,817,403 | 128,009,847 | 192,445 | 79 | CN0, CN2 | 0, 0.006 | esv3631107, esv3631108 | 1000G |
| CNP\_180 | chr12 | 129,230,099 | 129,233,221 | 3,123 | 5 | CN0 | 0.034 | esv3631149 | 1000G |
| CNP\_181 | chr12 | 130,008,238 | 130,256,986 | 248,749 | 136 | CN2 | 0.008 | esv3631178 | 1000G |
| CNP\_182 | chr12 | 130,402,334 | 130,582,861 | 180,528 | 85 | CN2 | 0.008 | esv3631192 | 1000G |
| CNP\_183 | chr12 | 130,860,279 | 131,092,219 | 231,941 | 160 | CN2 | 0.006 | esv3631200 | 1000G |
| CNP\_184 | chr13 | 34,135,729 | 34,144,821 | 9,093 | 4 | CN0 | 0.123 | esv3631741 | 1000G |
| CNP\_185 | chr13 | 57,715,850 | 57,826,089 | 110,240 | 9 | CN0 | 0.009 | esv3632213 | 1000G |
| CNP\_186 | chr13 | 69,244,692 | 69,268,758 | 24,067 | 10 | CN0 | 0.065 | esv3632602 | 1000G |
| CNP\_188 | chr13 | 76,107,422 | 76,121,306 | 13,885 | 6 | CN2 | 0 | esv3632749 | 1000G |
| CNP\_189 | chr13 | 114,545,142 | 114,548,662 | 3,521 | 4 | CN0, CN2 | 0.009, 0 | esv3633624, esv3633625 | 1000G |
| CNP\_191 | chr14 | 20,335,553 | 20,423,316 | 87,764 | 57 | CN2, CN3, CN4 | 0.3, 0.034, 0.013 | esv3633663, esv3633664, esv3633665 | 1000G |
| CNP\_192 | chr14 | 22,740,308 | 22,923,239 | 182,932 | 69 | CN0 | 0.005 | esv3633740 | 1000G |
| CNP\_193 | chr14 | 101,461,351 | 101,532,406 | 71,056 | 21 | CN2 | 0.066 | esv3635521 | 1000G |
| CNP\_194 | chr14 | 102,267,755 | 102,427,340 | 159,586 | 44 | CN0, CN2 | 0.003, 0.001 | esv3635536, esv3635537 | 1000G |
| CNP\_195 | chr14 | 105,415,167 | 105,417,877 | 2,711 | 10 | CN0 | 0.105 | esv3635631 | 1000G |
| CNP\_197 | chr15 | 22,343,965 | 22,372,302 | 28,338 | 12 | CN0, CN2, CN3, CN4, CN5, CN6 | 0.002, 0.322, 0.164, 0.006, 0.004, 0.003 | esv3635763, esv3635764, esv3635765, esv3635766, esv3635767, esv3635768 | 1000G |
| CNP\_198 | chr15 | 22,372,936 | 22,383,193 | 10,258 | 6 | CN0, CN2, CN3, CN4, CN5, CN6 | 0.086, 0.35, 0.015, 0.01, 0, 0 | esv3635770, esv3635771, esv3635772, esv3635773, esv3635774, esv3635775 | 1000G |
| CNP\_200 | chr15 | 34,721,236 | 34,836,826 | 115,591 | 10 | CN0, CN2 | 0.086, 0.004 | esv3636115, esv3636116 | 1000G |
| CNP\_201 | chr15 | 64,152,654 | 64,302,980 | 150,327 | 52 | CN0, CN2 | 0.001, 0.001 | esv3636720, esv3636721 | 1000G |
| CNP\_202 | chr15 | 86,064,810 | 86,182,627 | 117,818 | 95 | CN0, CN2 | 0.002, 0 | esv3637117, esv3637118 | 1000G |
| CNP\_203 | chr15 | 97,814,902 | 97,835,452 | 20,551 | 4 | CN0 | 0.052 | esv3637372 | 1000G |
| CNP\_204 | chr16 | 1,138,986 | 1,151,109 | 12,124 | 6 | CN2 | 0.027 | esv3637603 | 1000G |
| CNP\_205 | chr16 | 15,080,267 | 15,105,113 | 24,847 | 6 | CN2, CN3 | 0.808, 0.004 | esv3638015, esv3638016 | 1000G |
| CNP\_206 | chr16 | 19,945,551 | 19,967,585 | 22,035 | 5 | CN0 | 0.13 | esv3638126 | 1000G |
| CNP\_207 | chr16 | 22,623,100 | 22,783,757 | 160,658 | 36 | CN2, CN3, CN4 | 0.905, 0.008, 0 | esv3638212, esv3638213, esv3638214 | 1000G |
| CNP\_208 | chr16 | 28,614,507 | 28,626,916 | 12,410 | 8 | CN0, CN2, CN3, CN4, CN5 | 0.025, 0.155, 0.005, 0.005, 0 | esv3638338, esv3638339, esv3638340, esv3638341, esv3638342 | 1000G |
| CNP\_209 | chr16 | 34,457,085 | 34,712,399 | 255,315 | 13 | CN0, CN2 | 0, 0.1 | esv3638499, esv3638500 | 1000G |
| CNP\_210 | chr16 | 55,780,196 | 55,824,474 | 44,279 | 5 | CN0 | 0.015 | esv3638685 | 1000G |
| CNP\_211 | chr16 | 55,832,207 | 55,864,521 | 32,315 | 18 | CN0, CN2, CN3 | 0, 0.136, 0 | esv3638688, esv3638689, esv3638690 | 1000G |
| CNP\_212 | chr16 | 70,181,849 | 70,196,732 | 14,884 | 13 | CN0, CN2, CN3, CN4, CN5 | 0, 0.537, 0.023, 0.034, 0.003 | esv3638945, esv3638946, esv3638947, esv3638948, esv3638949 | 1000G |
| CNP\_213 | chr16 | 72,080,868 | 72,098,986 | 18,119 | 9 | CN0, CN2, CN3 | 0, 0, 0 | esv3638989, esv3638990, esv3638991 | 1000G |
| CNP\_214 | chr16 | 72,108,348 | 72,118,125 | 9,778 | 13 | CN2, CN3 | 0, 0 | esv3638998, esv3638999 | 1000G |
| CNP\_215 | chr17 | 10,886,864 | 10,895,981 | 9,118 | 5 | CN0 | 0.009 | esv3639945 | 1000G |
| CNP\_216 | chr17 | 14,224,374 | 14,483,419 | 259,046 | 142 | CN0, CN2 | 0, 0.003 | esv3640025, esv3640026 | 1000G |
| CNP\_217 | chr17 | 14,668,589 | 15,084,842 | 416,254 | 218 | CN0, CN2 | 0, 0.01 | esv3640045, esv3640046 | 1000G |
| CNP\_218 | chr17 | 15,043,705 | 15,058,719 | 15,015 | 9 | CN0 | 0.004 | esv3640059 | 1000G |
| CNP\_219 | chr17 | 34,436,099 | 34,482,872 | 46,774 | 16 | CN2, CN3, CN4 | 0.142, 0.019, 0.006 | esv3640465, esv3640466, esv3640467 | 1000G |
| CNP\_220 | chr17 | 39,506,753 | 39,525,903 | 19,151 | 9 | CN0 | 0 | esv3640584 | 1000G |
| CNP\_221 | chr17 | 39,531,703 | 39,539,741 | 8,039 | 12 | CN2 | 0.025 | esv3640585 | 1000G |
| CNP\_222 | chr17 | 44,165,338 | 44,211,686 | 46,349 | 8 | CN2, CN3 | 0.188, 0.005 | esv3640677, esv3640678 | 1000G |
| CNP\_223 | chr17 | 44,230,893 | 44,262,697 | 31,805 | 27 | CN2, CN3 | 0.41, 0.005 | esv3640680, esv3640681 | 1000G |
| CNP\_224 | chr17 | 54,160,158 | 54,172,953 | 12,796 | 9 | CN0 | 0.053 | esv3640845 | 1000G |
| CNP\_225 | chr17 | 54,161,689 | 54,167,620 | 5,932 | 5 | CN0 | 0.053 | esv3640846 | 1000G |
| CNP\_226 | chr17 | 79,100,329 | 79,110,432 | 10,104 | 5 | CN2 | 0.026 | esv3641392 | 1000G |
| CNP\_227 | chr18 | 2,659,472 | 2,734,289 | 74,818 | 20 | CN0, CN2 | 0.002, 0.003 | esv3641565, esv3641566 | 1000G |
| CNP\_228 | chr18 | 3,200,017 | 3,415,245 | 215,229 | 95 | CN0, CN2 | 0.007, 0 | esv3641584, esv3641585 | 1000G |
| CNP\_229 | chr18 | 66,745,584 | 66,756,995 | 11,412 | 9 | CN0 | 0.028 | esv3642958 | 1000G |
| CNP\_230 | chr19 | 20,595,835 | 20,717,950 | 122,116 | 14 | CN0 | 0.06 | esv3643884 | 1000G |
| CNP\_232 | chr19 | 21,055,692 | 21,098,732 | 43,041 | 6 | CN2, CN3 | 0.116, 0.003 | esv3643908, esv3643909 | 1000G |
| CNP\_233 | chr19 | 28,246,833 | 28,474,547 | 227,715 | 46 | CN2, CN3 | 0.002, 0 | esv3644080, esv3644081 | 1000G |
| CNP\_234 | chr19 | 35,661,065 | 35,665,796 | 4,732 | 4 | CN0 | 0.075 | esv3644227 | 1000G |
| CNP\_235 | chr19 | 40,519,935 | 40,541,914 | 21,980 | 27 | NA | NA |  | Panc4 |
| CNP\_236 | chr19 | 41,352,463 | 41,383,107 | 30,645 | 12 | CN0 | 0.023 | esv3644361 | 1000G |
| CNP\_237 | chr19 | 41,354,052 | 41,373,538 | 19,487 | 10 | CN0, CN2 | 0.028, 0.009 | esv3644362, esv3644363 | 1000G |
| CNP\_239 | chr19 | 41,361,249 | 41,365,764 | 4,516 | 5 | CN0 | 0.022 | esv3644364 | 1000G |
| CNP\_241 | chr19 | 43,423,930 | 43,451,047 | 27,118 | 12 | CN0, CN2 | 0.025, 0.003 | esv3644425, esv3644426 | 1000G |
| CNP\_242 | chr19 | 43,505,313 | 43,547,211 | 41,899 | 10 | CN0, CN2 | 0.033, 0.003 | esv3644435, esv3644436 | 1000G |
| CNP\_243 | chr19 | 46,622,653 | 46,628,256 | 5,604 | 5 | CN0 | 0.018 | esv3644520 | 1000G |
| CNP\_244 | chr19 | 53,520,294 | 53,545,025 | 24,732 | 9 | CN0, CN2, CN3 | 0, 0.082, 0.018 | esv3644732, esv3644733, esv3644734 | 1000G |
| CNP\_245 | chr19 | 54,727,998 | 54,746,910 | 18,913 | 12 | CN0, CN2, CN3 | 0.012, 0.246, 0.006 | esv3644786, esv3644787, esv3644788 | 1000G |
| CNP\_246 | chr19 | 54,800,423 | 54,807,689 | 7,267 | 8 | CN0 | 0.178 | esv3644795 | 1000G |
| CNP\_247 | chr19 | 55,264,338 | 55,367,571 | 103,234 | 23 | CN0 | 0.025 | esv3644815 | 1000G |
| CNP\_248 | chr19 | 55,338,842 | 55,355,281 | 16,440 | 5 | CN0, CN2 | 0.227, 0.001 | esv3644818, esv3644819 | 1000G |
| CNP\_249 | chr20 | 1,557,325 | 1,559,946 | 2,622 | 4 | CN0 | 0.141 | esv3644982 | 1000G |
| CNP\_250 | chr20 | 58,436,963 | 58,496,439 | 59,477 | 49 | NA | NA |  | Panc4 |
| CNP\_251 | chr21 | 10,824,040 | 10,942,992 | 118,953 | 19 | NA | NA |  | Panc4 |
| CNP\_252 | chr21 | 23,654,294 | 23,666,590 | 12,297 | 4 | CN0 | 0.064 | esv3646598 | 1000G |
| CNP\_253 | chr21 | 28,195,256 | 28,201,949 | 6,694 | 4 | CN0 | 0 | esv3646741 | 1000G |
| CNP\_254 | chr22 | 23,027,159 | 23,088,641 | 61,483 | 4 | CN0 | 0.023 | esv3647376 | 1000G |
| CNP\_255 | chr22 | 25,659,945 | 25,710,725 | 50,781 | 9 | CN0, CN2, CN3 | 0.015, 0.021, 0 | esv3647453, esv3647454, esv3647455 | 1000G |
| CNP\_257 | chr22 | 25,703,077 | 25,784,925 | 81,849 | 37 | CN0, CN2, CN3, CN4 | 0.015, 0.02, 0, 0 | esv3647457, esv3647458, esv3647459, esv3647460 | 1000G |
| CNP\_258 | chr22 | 25,825,377 | 25,844,076 | 18,700 | 5 | CN0, CN2, CN3, CN4 | 0.015, 0.019, 0.001, 0 | esv3647470, esv3647471, esv3647472, esv3647473 | 1000G |
| CNP\_259 | chr22 | 25,846,677 | 25,921,876 | 75,200 | 19 | CN0, CN2, CN3 | 0.015, 0.02, 0 | esv3647474, esv3647475, esv3647476 | 1000G |
| CNP\_260 | chr22 | 39,357,694 | 39,388,574 | 30,881 | 6 | CN0 | 0.069 | esv3647746 | 1000G |
| CNP\_261 | chr22 | 42,523,949 | 42,533,891 | 9,943 | 8 | CN0, CN2, CN3, CN4 | 0.025, 0.084, 0.001, 0.001 | esv3647809, esv3647810, esv3647811, esv3647812 | 1000G |
| CNP\_262 | chr22 | 42,906,659 | 42,934,146 | 27,488 | 13 | CN0, CN2, CN3 | 0.001, 0.122, 0.005 | esv3647831, esv3647832, esv3647833 | 1000G |
| CNP\_263 | chr22 | 49,401,061 | 49,500,765 | 99,705 | 47 | CN2 | 0.006 | esv3648032 | 1000G |

### Table S2: Posterior probabilities of association

```
..............
```

| CNP id | Location | Gene | Copy number genotypes | Probability of association | CNVs pancreatic cancer | CNVs healthy controls |
| --- | --- | --- | --- | --- | --- | --- |
| CNP\_003 | chr1:12,856,327-12,893,193 | PRAMEF1 | 1,2,3 | 0.037 | 226 | 208 |
| CNP\_004 | chr1:12,877,893-12,933,272 | PRAMEF2 | 0,1,2 | 0.002 | 311 | 274 |
| CNP\_005 | chr1:12,893,536-12,948,772 | PRAMEF2 | 0,1,2 | 0.004 | 312 | 276 |
| CNP\_006 | chr1:12,896,438-12,928,826 | PRAMEF2 | 0,1,2 | 0.002 | 293 | 262 |
| CNP\_007 | chr1:12,901,370-12,921,250 | PRAMEF2 | 0,1,2,3 | 0.002 | 424 | 384 |
| CNP\_008 | chr1:12,911,200-12,921,311 | PRAMEF2 | 0,1,2 | 0.002 | 406 | 333 |
| CNP\_014 | chr1:25,592,642-25,661,222 | RHD | 1,2 | 0.010 | 547 | 574 |
| CNP\_016 | chr1:104,102,743-104,136,676 | RNPC3 | 2,3 | 0.009 | 451 | 369 |
| CNP\_017 | chr1:104,107,068-104,211,046 | RNPC3 | 2,3 | 0.007 | 505 | 394 |
| CNP\_018 | chr1:109,366,972-109,372,000 | AKNAD1 | 0,1,2 | 0.001 | 93 | 77 |
| CNP\_019 | chr1:110,225,019-110,245,280 | GSTM2 | 1,2 | 0.000 | 2040 | 1886 |
| CNP\_020 | chr1:110,230,075-110,241,247 | GSTM2 | 1,2 | 0.001 | 2104 | 1925 |
| CNP\_022 | chr1:174,796,517-174,801,833 | RABGAP1L | 0,1,2 | 0.001 | 874 | 863 |
| CNP\_023 | chr1:196,801,025-196,892,322 | CFHR1 | 0,1,2,3 | 0.023 | 79 | 91 |
| CNP\_026 | chr2:41,238,373-41,250,303 | NA | 0,1,2 | 0.002 | 592 | 533 |
| CNP\_027 | chr2:49,532,573-49,542,423 | NA | 0,1,2 | 0.005 | 336 | 322 |
| CNP\_028 | chr2:51,926,595-51,927,817 | NA | 0,1,2 | 0.002 | 779 | 737 |
| CNP\_029 | chr2:90,010,895-90,248,037 | NA | 0,1,2,3 | 0.002 | 292 | 249 |
| CNP\_030 | chr2:90,066,423-90,120,854 | NA | 0,1,2,3 | 0.002 | 292 | 254 |
| CNP\_031 | chr2:90,212,408-90,249,248 | NA | 0,1,2 | 0.013 | 262 | 201 |
| CNP\_033 | chr2:195,979,320-195,984,295 | NA | 0,1,2 | 0.007 | 535 | 423 |
| CNP\_034 | chr2:208,351,101-208,359,313 | NA | 0,1,2 | 0.001 | 1260 | 1103 |
| CNP\_035 | chr2:213,183,956-213,192,007 | ERBB4 | 0,1,2 | 0.002 | 522 | 468 |
| CNP\_036 | chr2:242,917,734-243,044,147 | LINC01237 | 0,1,2 | 0.002 | 316 | 310 |
| CNP\_037 | chr3:37,978,345-37,986,932 | CTDSPL | 0,1,2 | 0.002 | 607 | 599 |
| CNP\_038 | chr3:65,188,826-65,214,723 | NA | 0,1,2 | 0.001 | 545 | 518 |
| CNP\_039 | chr3:65,191,847-65,214,533 | NA | 0,1,2 | 0.001 | 545 | 516 |
| CNP\_040 | chr3:75,428,675-75,703,846 | MIR1324 | 0,1,2 | 0.002 | 485 | 470 |
| CNP\_041 | chr3:148,963,333-148,969,710 | NA | 0,1,2 | 0.001 | 316 | 289 |
| CNP\_042 | chr3:162,127,885-162,143,633 | NA | 0,1,2 | 0.011 | 463 | 392 |
| CNP\_044 | chr3:173,239,453-173,289,281 | NLGN1 | 2,3 | 0.006 | 177 | 165 |
| CNP\_045 | chr3:191,064,690-191,071,659 | CCDC50 | 0,1,2 | 0.001 | 1718 | 1533 |
| CNP\_047 | chr3:193,135,464-193,142,938 | ATP13A4 | 0,1,2 | 0.023 | 172 | 148 |
| CNP\_053 | chr4:10,392,397-10,402,099 | NA | 0,1,2 | 0.000 | 1732 | 1602 |
| CNP\_055 | chr4:64,133,825-64,153,198 | NA | 0,1,2 | 0.002 | 658 | 614 |
| CNP\_056 | chr4:69,375,591-69,491,543 | UGT2B17 | 0,2 | 0.000 | 429 | 407 |
| CNP\_059 | chr4:122,282,235-122,289,973 | QRFPR | 0,1,2 | 0.013 | 438 | 415 |
| CNP\_061 | chr4:152,790,167-152,794,773 | NA | 0,1,2 | 0.001 | 496 | 470 |
| CNP\_062 | chr4:161,059,762-161,071,059 | NA | 0,1,2 | 0.004 | 225 | 192 |
| CNP\_063 | chr5:788,646-821,992 | ZDHHC11 | 0,1,2 | 0.002 | 315 | 286 |
| CNP\_064 | chr5:814,446-825,367 | ZDHHC11 | 0,1,2 | 0.014 | 242 | 236 |
| CNP\_065 | chr5:32,107,381-32,159,517 | PDZD2 | 2,3 | 0.004 | 205 | 182 |
| CNP\_067 | chr5:57,320,371-57,338,133 | NA | 1,2 | 0.001 | 102 | 75 |
| CNP\_068 | chr5:70,305,696-70,308,554 | NAIP | 0,1,2,3 | 0.006 | 741 | 757 |
| CNP\_069 | chr5:70,306,516-70,313,302 | NAIP | 0,1,2,3 | 0.003 | 670 | 665 |
| CNP\_070 | chr5:97,048,466-97,099,320 | NA | 0,1,2 | 0.004 | 261 | 238 |
| CNP\_072 | chr5:140,222,274-140,239,347 | PCDHA1 | 0,1,2 | 0.015 | 697 | 713 |
| CNP\_073 | chr5:140,223,185-140,238,124 | PCDHA1 | 0,1,2 | 0.032 | 696 | 710 |
| CNP\_078 | chr5:151,514,809-151,518,869 | CTB-12O2.1 | 0,1,2 | 0.004 | 1143 | 1010 |
| CNP\_079 | chr5:155,475,886-155,488,649 | NA | 0,1,2 | 0.004 | 1276 | 1109 |
| CNP\_081 | chr6:29,851,172-29,904,515 | HLA-H | 0,1,2 | 0.001 | 838 | 748 |
| CNP\_082 | chr6:29,859,708-29,888,317 | HLA-H | 0,1,2 | 0.002 | 839 | 750 |
| CNP\_083 | chr6:30,994,015-30,995,083 | MUC22 | 0,1,2 | 0.000 | 857 | 802 |
| CNP\_085 | chr6:31,219,515-31,229,793 | HLA-C | 1,2 | 0.039 | 131 | 104 |
| CNP\_086 | chr6:31,275,976-31,279,146 | NA | 0,1,2 | 0.003 | 652 | 564 |
| CNP\_087 | chr6:31,337,847-31,341,985 | NA | 0,1,2 | 0.013 | 121 | 90 |
| CNP\_091 | chr6:67,009,228-67,049,033 | NA | 0,1,2 | 0.001 | 595 | 583 |
| CNP\_092 | chr6:67,017,494-67,047,294 | NA | 0,1,2 | 0.001 | 594 | 581 |
| CNP\_094 | chr6:78,892,808-79,053,430 | NA | 0,2 | 0.000 | 1775 | 1633 |
| CNP\_095 | chr6:78,967,097-79,036,275 | NA | 0,1,2 | 0.000 | 1695 | 1603 |
| CNP\_096 | chr6:78,972,930-79,029,367 | NA | 0,1,2 | 0.000 | 1693 | 1607 |
| CNP\_097 | chr6:81,283,720-81,293,577 | NA | 0,1,2 | 0.007 | 447 | 366 |
| CNP\_099 | chr6:168,364,813-168,593,956 | MLLT4 | 2,3 | 0.011 | 105 | 96 |
| CNP\_100 | chr7:39,119-75,937 | NA | 2,3 | 0.067 | 170 | 187 |
| CNP\_101 | chr7:44,935-68,920 | NA | 2,3 | 0.007 | 190 | 188 |
| CNP\_105 | chr7:76,417,751-76,615,349 | DTX2P1-UPK3BP1-PMS2P11 | 1,2,3 | 0.012 | 91 | 108 |
| CNP\_109 | chr7:141,765,307-141,792,849 | MGAM | 0,1,2 | 0.000 | 1341 | 1257 |
| CNP\_110 | chr7:142,475,484-142,486,103 | PRSS3P2 | 0,1,2 | 0.001 | 457 | 398 |
| CNP\_111 | chr7:142,827,954-142,881,540 | PIP | 0,1,2 | 0.024 | 90 | 96 |
| CNP\_113 | chr8:3,786,311-3,790,617 | CSMD1 | 0,1,2 | 0.002 | 421 | 414 |
| CNP\_115 | chr8:5,595,438-5,605,648 | NA | 0,1,2 | 0.001 | 593 | 541 |
| CNP\_116 | chr8:5,599,399-5,605,087 | NA | 0,1,2 | 0.000 | 591 | 540 |
| CNP\_121 | chr8:15,401,664-15,414,791 | TUSC3 | 0,1,2 | 0.073 | 211 | 220 |
| CNP\_122 | chr8:16,262,082-16,274,699 | NA | 0,1,2 | 0.015 | 353 | 292 |
| CNP\_124 | chr8:40,182,839-40,189,804 | NA | 0,1,2 | 0.002 | 407 | 407 |
| CNP\_125 | chr8:85,260,961-85,269,169 | RALYL | 0,1,2 | 0.005 | 248 | 265 |
| CNP\_128 | chr8:137,682,484-137,857,327 | NA | 0,1,2 | 0.183 | 147 | 174 |
| CNP\_135 | chr10:20,849,543-20,860,166 | MIR4675 | 0,1,2 | 0.000 | 1257 | 1175 |
| CNP\_136 | chr10:47,531,022-47,599,037 | ANTXRLP1 | 1,2,3 | 0.003 | 453 | 395 |
| CNP\_137 | chr10:47,543,322-47,703,869 | ANTXRL | 2,3 | 0.009 | 386 | 351 |
| CNP\_139 | chr10:47,645,964-47,697,149 | ANTXRL | 1,2,3 | 0.024 | 416 | 386 |
| CNP\_140 | chr10:58,512,456-58,527,377 | NA | 0,1,2 | 0.002 | 363 | 325 |
| CNP\_141 | chr10:68,078,481-68,114,481 | CTNNA3 | 0,1,2 | 0.008 | 92 | 80 |
| CNP\_146 | chr10:135,256,762-135,369,532 | SCART1 | 2,3 | 0.013 | 172 | 141 |
| CNP\_147 | chr10:135,348,035-135,378,260 | CYP2E1 | 2,3 | 0.002 | 186 | 145 |
| CNP\_149 | chr11:3,238,736-3,244,087 | MRGPRG-AS1 | 0,1,2 | 0.003 | 375 | 335 |
| CNP\_152 | chr11:7,812,280-7,832,579 | OR5P2 | 0,1,2 | 0.009 | 305 | 257 |
| CNP\_153 | chr11:18,941,736-18,963,997 | MRGPRX1 | 0,1,2 | 0.010 | 185 | 188 |
| CNP\_155 | chr11:55,368,372-55,426,969 | OR4P4 | 0,1,2 | 0.000 | 1785 | 1662 |
| CNP\_156 | chr11:55,370,325-55,427,700 | OR4P4 | 0,1,2 | 0.000 | 1789 | 1667 |
| CNP\_158 | chr11:81,500,537-81,521,332 | NA | 0,1,2 | 0.001 | 649 | 594 |
| CNP\_159 | chr11:86,304,229-86,306,559 | ME3 | 0,1,2 | 0.002 | 762 | 649 |
| CNP\_163 | chr12:8,000,912-8,114,429 | SLC2A14 | 1,2,3 | 0.022 | 195 | 172 |
| CNP\_166 | chr12:11,222,191-11,249,671 | PRH1-PRR4 | 0,1,2 | 0.007 | 845 | 816 |
| CNP\_167 | chr12:27,648,174-27,655,202 | SMCO2 | 0,1,2 | 0.004 | 214 | 172 |
| CNP\_168 | chr12:31,266,287-31,407,303 | DDX11 | 2,3 | 0.011 | 274 | 294 |
| CNP\_169 | chr12:33,296,520-33,307,372 | NA | 0,1,2 | 0.001 | 1548 | 1386 |
| CNP\_171 | chr12:53,086,320-53,087,693 | KRT77 | 0,1,2 | 0.001 | 522 | 514 |
| CNP\_173 | chr12:70,679,288-70,682,287 | CNOT2 | 0,1,2 | 0.002 | 401 | 353 |
| CNP\_174 | chr12:70,872,239-70,878,209 | NA | 0,1,2 | 0.000 | 1266 | 1156 |
| CNP\_180 | chr12:129,230,099-129,233,221 | NA | 0,1,2 | 0.017 | 281 | 244 |
| CNP\_184 | chr13:34,135,729-34,144,821 | STARD13 | 0,1,2 | 0.002 | 718 | 629 |
| CNP\_186 | chr13:69,244,692-69,268,758 | NA | 0,1,2 | 0.021 | 430 | 448 |
| CNP\_187 | chr13:69,247,022-69,267,981 | NA | 0,1,2 | 0.026 | 430 | 448 |
| CNP\_190 | chr14:19,847,604-20,404,736 | BMS1P18 | 2,3 | 0.019 | 1287 | 1179 |
| CNP\_191 | chr14:20,335,553-20,423,316 | OR4K2 | 2,3 | 0.004 | 1276 | 1167 |
| CNP\_198 | chr15:22,372,936-22,383,193 | LOC101927079 | 0,1,2 | 0.004 | 121 | 119 |
| CNP\_199 | chr15:34,718,594-34,807,851 | GOLGA8A | 0,1,2 | 0.032 | 669 | 534 |
| CNP\_200 | chr15:34,721,236-34,836,826 | GOLGA8A | 0,1,2 | 0.038 | 668 | 532 |
| CNP\_203 | chr15:97,814,902-97,835,452 | NA | 0,1,2 | 0.001 | 402 | 361 |
| CNP\_206 | chr16:19,945,551-19,967,585 | NA | 0,1,2 | 0.001 | 1003 | 942 |
| CNP\_208 | chr16:28,614,507-28,626,916 | SULT1A2 | 0,1,2 | 0.007 | 174 | 152 |
| CNP\_209 | chr16:34,457,085-34,712,399 | LINC01566 | 2,3 | 0.007 | 625 | 634 |
| CNP\_211 | chr16:55,832,207-55,864,521 | CES1 | 1,2 | 0.005 | 3364 | 3083 |
| CNP\_219 | chr17:34,436,099-34,482,872 | CCL4 | 2,3 | 0.001 | 864 | 759 |
| CNP\_221 | chr17:39,531,703-39,539,741 | KRT33B | 2,3 | 0.010 | 112 | 87 |
| CNP\_222 | chr17:44,165,338-44,211,686 | KANSL1 | 2,3 | 0.023 | 1168 | 1152 |
| CNP\_223 | chr17:44,230,893-44,262,697 | KANSL1-AS1 | 2,3 | 0.002 | 2370 | 2183 |
| CNP\_224 | chr17:54,160,158-54,172,953 | NA | 0,1,2 | 0.003 | 304 | 265 |
| CNP\_225 | chr17:54,161,689-54,167,620 | NA | 0,1,2 | 0.001 | 298 | 261 |
| CNP\_229 | chr18:66,745,584-66,756,995 | NA | 0,1,2 | 0.001 | 265 | 268 |
| CNP\_230 | chr19:20,595,835-20,717,950 | ZNF826P | 0,1,2 | 0.003 | 484 | 409 |
| CNP\_231 | chr19:20,621,828-20,715,228 | ZNF737 | 0,1,2 | 0.002 | 484 | 409 |
| CNP\_234 | chr19:35,661,065-35,665,796 | FXYD5 | 0,1,2 | 0.010 | 495 | 509 |
| CNP\_236 | chr19:41,352,463-41,383,107 | CYP2A6 | 0,1,2,3 | 0.010 | 276 | 236 |
| CNP\_237 | chr19:41,354,052-41,373,538 | CYP2A6 | 0,1,2,3 | 0.006 | 280 | 245 |
| CNP\_238 | chr19:41,355,999-41,386,033 | CYP2A6 | 0,1,2 | 0.008 | 210 | 178 |
| CNP\_239 | chr19:41,361,249-41,365,764 | CYP2A6 | 0,1,2 | 0.030 | 181 | 151 |
| CNP\_240 | chr19:43,328,006-43,702,355 | LOC100289650 | 0,1,2,3 | 0.214 | 126 | 143 |
| CNP\_241 | chr19:43,423,930-43,451,047 | PSG6 | 0,1,2 | 0.029 | 90 | 106 |
| CNP\_242 | chr19:43,505,313-43,547,211 | PSG11 | 0,1,2 | 0.004 | 162 | 176 |
| CNP\_243 | chr19:46,622,653-46,628,256 | IGFL3 | 0,1,2 | 0.008 | 97 | 98 |
| CNP\_244 | chr19:53,520,294-53,545,025 | ERVV-1 | 2,3 | 0.008 | 730 | 699 |
| CNP\_245 | chr19:54,727,998-54,746,910 | LILRB3 | 0,1,2,3 | 0.005 | 316 | 265 |
| CNP\_246 | chr19:54,800,423-54,807,689 | LILRA3 | 1,2 | 0.005 | 159 | 158 |
| CNP\_248 | chr19:55,338,842-55,355,281 | KIR3DL1 | 0,1,2 | 0.008 | 137 | 129 |
| CNP\_249 | chr20:1,557,325-1,559,946 | SIRPB1 | 0,1,2 | 0.001 | 961 | 886 |
| CNP\_252 | chr21:23,654,294-23,666,590 | NA | 0,1,2 | 0.005 | 540 | 465 |
| CNP\_254 | chr22:23,027,159-23,088,641 | NA | 0,1,2 | 0.074 | 340 | 244 |
| CNP\_255 | chr22:25,659,945-25,710,725 | IGLL3P | 1,2,3 | 0.015 | 210 | 166 |
| CNP\_256 | chr22:25,668,730-25,914,593 | IGLL3P | 1,2,3 | 0.003 | 254 | 202 |
| CNP\_257 | chr22:25,703,077-25,784,925 | IGLL3P | 1,2,3 | 0.004 | 233 | 174 |
| CNP\_259 | chr22:25,846,677-25,921,876 | CRYBB2P1 | 1,2,3 | 0.002 | 256 | 200 |
| CNP\_260 | chr22:39,357,694-39,388,574 | APOBEC3A | 1,2 | 0.006 | 21 | 35 |
| CNP\_262 | chr22:42,906,659-42,934,146 | SERHL | 2,3 | 0.003 | 722 | 670 |

### Table S3: Hyperparameters for Bayesian mixture model

| Hyperparameter | Value |
| --- | --- |
| \(\mu\_0\) | 0 |
| \(\tau^2\) | 0.4 |
| \(\eta\_0\) | 32 |
| \(m^2\) | 0.5 |
| \(\alpha\) | \(\mathbf{1}\) |
| \(\beta\) | 0.1 |
| a | 1.8 |
| b | 6 |
| df | 100 |

### Table S4: Allelic copy numbers

| Total copy number | Allelic copy number |
| --- | --- |
| 0 | NULL |
| 1 | A, B |
| 2 | AA, AB, BB |
| 3 | AAA, AAB, ABB, BBB |
| 4 | AAAA, AAAB, AABB, ABBB, BBB |

### Table S5: Shape and scale parameters of beta distribution for the different allelic copy numbers

| Allelic copy number | \(\psi\) |
| --- | --- |
| NULL | 1, 1 |
| B, BB, BBB, BBBB | 10, 1 |
| A, AA, AAA, AAAA | 1, 10 |
| AB, AABB | 30, 30 |
| AAB | 10, 20 |
| ABB | 20, 10 |
| AAAB | 30, 10 |
| ABBB | 10, 30 |

  

Session information

```
R version 3.6.2 (2019-12-12)
Platform: x86_64-apple-darwin19.2.0/x86_64 (64-bit)
Running under: macOS Catalina 10.15.2

Matrix products: default
BLAS:   /Users/rscharpf/Rversions/R-3.6.2/lib/x86_64/libRblas.dylib
LAPACK: /Users/rscharpf/Rversions/R-3.6.2/lib/x86_64/libRlapack.dylib

locale:
[1] en_US.UTF-8/en_US.UTF-8/en_US.UTF-8/C/en_US.UTF-8/en_US.UTF-8

attached base packages:
[1] parallel  stats4    stats     graphics  grDevices utils     datasets 
[8] methods   base     

other attached packages:
 [1] regression.models_0.0.1     panc.funs_0.0.2            
 [3] panc.data_0.0.2             magrittr_1.5               
 [5] CNPBayes_1.15.5             SummarizedExperiment_1.16.1
 [7] DelayedArray_0.12.2         BiocParallel_1.20.1        
 [9] matrixStats_0.55.0          Biobase_2.46.0             
[11] GenomicRanges_1.38.0        GenomeInfoDb_1.22.0        
[13] IRanges_2.20.1              S4Vectors_0.24.1           
[15] BiocGenerics_0.32.0         kableExtra_1.1.0           
[17] forcats_0.4.0               stringr_1.4.0              
[19] dplyr_0.8.3                 purrr_0.3.3                
[21] readr_1.3.1                 tidyr_1.0.0                
[23] tibble_2.1.3                ggplot2_3.2.1              
[25] tidyverse_1.3.0             knitr_1.26                 
[27] workflowr_1.6.0            

loaded via a namespace (and not attached):
 [1] nlme_3.1-143           bitops_1.0-6           fs_1.3.1              
 [4] lubridate_1.7.4        webshot_0.5.2          RColorBrewer_1.1-2    
 [7] httr_1.4.1             rprojroot_1.3-2        tools_3.6.2           
[10] backports_1.1.5        R6_2.4.1               DBI_1.1.0             
[13] lazyeval_0.2.2         colorspace_1.4-1       withr_2.1.2           
[16] tidyselect_0.2.5       compiler_3.6.2         git2r_0.26.1          
[19] cli_2.0.0              rvest_0.3.5            xml2_1.2.2            
[22] scales_1.1.0           mvtnorm_1.0-11         digest_0.6.23         
[25] rmarkdown_2.0          XVector_0.26.0         pkgconfig_2.0.3       
[28] htmltools_0.4.0        dbplyr_1.4.2           highr_0.8             
[31] rlang_0.4.2            readxl_1.3.1           rstudioapi_0.10       
[34] generics_0.0.2         combinat_0.0-8         jsonlite_1.6          
[37] gtools_3.8.1           RCurl_1.95-4.12        GenomeInfoDbData_1.2.2
[40] Matrix_1.2-18          Rcpp_1.0.3             munsell_0.5.0         
[43] fansi_0.4.0            lifecycle_0.1.0        stringi_1.4.3         
[46] yaml_2.2.0             zlibbioc_1.32.0        grid_3.6.2            
[49] promises_1.1.0         crayon_1.3.4           lattice_0.20-38       
[52] haven_2.2.0            hms_0.5.2              zeallot_0.1.0         
[55] pillar_1.4.3           reprex_0.3.0           glue_1.3.1            
[58] evaluate_0.14          modelr_0.1.5           vctrs_0.2.1           
[61] httpuv_1.5.2           cellranger_1.1.0       gtable_0.3.0          
[64] assertthat_0.2.1       xfun_0.11              broom_0.5.3           
[67] coda_0.19-3            later_1.0.0            viridisLite_0.3.0     
[70] ellipsis_0.3.0
```
